# Analyzing long-read CRISPR experiments with CRISPRLungo

**DOI:** 10.1101/2025.10.21.683786

**Authors:** Gue-Ho Hwang, Benjamin Vyshedskiy, Timothy Barry, Jing Zeng, John P. Manis, Akiko Shimamura, Daniel E. Bauer, Luca Pinello

## Abstract

Long-read sequencing can characterize complex genome editing-induced DNA sequence changes such as large deletions, insertions, and inversions that are difficult to detect using short-read sequencing. However, PCR amplification and sequencing errors complicate accurate variant detection, and existing analysis tools are not optimized for gene editing specific allelic outcomes. Here we present CRISPRLungo, a computational pipeline specifically designed for long-read amplicon sequencing of gene edited samples. CRISPRLungo incorporates unique molecular identifier (UMI)-based error correction and statistical filtering to distinguish true editing events from background noise, enabling robust detection of small indels and structural variants. Through systematic benchmarking using simulated datasets, we demonstrate that CRISPRLungo outperforms existing approaches in both accuracy and read recovery. CRISPRLungo supports both Oxford Nanopore and PacBio platforms and identify previously undetected structural variant edits such as inversions in published CRISPR datasets. To demonstrate allele-specific edit quantification, we applied CRISPRLungo to analyze edited primary cells from a patient with harboring compound heterozygous *SBDS* mutations, accurately quantifying *SBDS* editing outcomes despite contaminating reads from the homologous *SBDSP1* pseudogene. To maximize accessibility, we developed a fully client-side web application requiring no installation, making advanced long-read analysis accessible to researchers regardless of computational expertise. CRISPRLungo is freely available at https://github.com/pinellolab/CRISPRLungo with a user-friendly web interface available at https://pinellolab.github.io/CRISPRLungo.

## INTRODUCTION

CRISPR genome editing systems have revolutionized our ability to precisely modify DNA, offering programmable nucleases that induce double-stranded DNA breaks (DSBs) to introduce targeted genetic alterations^1, 2^. Beyond traditional nucleases, Cas proteins have been engineered into base editors (BEs)^3, 4^, which couple deaminases to Cas9 nickase (nCas9)^5^, and prime editors (PEs)^6, 7^, which combine nCas9 with reverse transcriptase. Recent innovations including dual flap prime editing^8, 9^ and recombinase systems^10^, have further expanded editing capabilities, enabling precise integration and modification of larger DNA sequences. These advances are accelerating CRISPR applications across disease modeling, agriculture, and therapeutic development^11-13^.

As CRISPR technologies mature, accurately analyzing both on-target and off-target editing outcomes becomes essential^14-16^. While early analyses focused primarily on detecting small insertions and deletions (indels) using short-read sequencing, recent studies reveal that CRISPR-Cas editing frequently induces structural variant outcomes including large deletions and rearrangements^17^. Even base and prime editors, designed to avoid DSBs through nCas9, can generate large deletions under certain conditions^18^. To characterize these events, two principal approaches have emerged: (i) single-anchor amplicon sequencing methods, such as UDiTaS^19^ and PEM-seq^20^, which detect unidirectional large deletions and translocations, and (ii) long-read amplicon sequencing platforms, including Oxford Nanopore and PacBio, which enable bidirectional detection of deletions, insertions, inversions, and duplications^17, 21^. While single-anchor methods represented an advance, they remain semiquantitative due to amplification biases and cannot detect all structural variants, particularly inversions and complex rearrangements that require bidirectional sequence information. While single-anchor methods improve detection of some structural variants, they are limited by short-read length constraints and may miss complex rearrangements. Long-read approaches overcome these length limitations but suffer from sequencing and PCR-induced artifacts that can obscure true editing events^22, 23^.

Unique molecular identifier (UMI) long-read sequencing strategies have been developed to improve accuracy by collapsing reads into error-corrected consensus sequences^24, 25^, and these methods are becoming increasingly adopted, necessitating specialized analytical frameworks. While several tools have emerged, each has significant limitations. For example, Oxford Nanopore’s *pipeline-umi-amplicon* enables UMI consensus generation but lacks CRISPR-specific mutational classification. Another study proposed instead a two-part workflow combining *longread_umi*^*25*^ and *LV_caller* (Large Variants caller)^24^, to detect large-scale mutations. However, this pipeline (hereafter referred to as the “*longread_umi + LV_caller*”) does not capture common CRISPR editing outcomes, such as small indels or inversions. In addition, reliance on command-line execution for both approaches creates accessibility barriers for researchers without bioinformatics expertise, limiting adoption in many research laboratories.

Here we present CRISPRLungo, a comprehensive analysis pipeline specifically designed for long-read amplicon sequencing of CRISPR-edited samples. CRISPRLungo accurately detects CRISPR-induced mutations, including small indels, large deletions, insertions, and critically, inversions, a mutation type overlooked by existing tools. The pipeline performs integrated end-to-end analysis, from UMI consensus generation to mutational classification, and incorporates statistical procedures to distinguish true mutation signatures from background noise arising from PCR amplification and long-read sequencing. Using systematic benchmarking on simulated datasets, we demonstrate that CRISPRLungo surpasses existing pipelines in both accuracy and read recovery. Importantly, validation with orthogonal droplet digital PCR (ddPCR) confirms quantification accuracy.

To demonstrate clinical utility in a challenging scenario, we applied CRISPRLungo to analyze allele-specific editing in primary cells from a patient with Schwachman-Diamond syndrome, accurately quantifying edited outcomes at the target *SBDS* locus while distinguishing contaminating reads from the highly homologous *SBDSP1* pseudogene^26^. Finally, to maximize accessibility and democratize access to advanced long-read analysis, we developed a fully client-side web application requiring no installation or bioinformatics expertise.

## RESULTS

### CRISPRLungo: a comprehensive pipeline for CRISPR mutation analysis

We developed CRISPRLungo as a modular pipeline that addresses key challenges in long-read sequencing analysis of gene-edited samples. CRISPRLungo consists of five integrated modules (**Fig. 1a**). i) Preprocessing: Minimap2^27^ removes low-quality reads and separates chimeric reads. ii) Consensus generation: When UMI sequences are present, CRISPRLungo performs error correction through UMI clustering and consensus generation (see **Methods** for optimization details). iii) Variant calling: Aligned reads are analyzed to identify mutation types, including small indels, large deletions, insertions, duplications, and inversions. iv) Statistical filtering: When control samples are provided, statistical comparison between treated and control datasets identifies true editing patterns and remove background signals. v) Visualization: Final mutation events are quantified and visualized through several informative plots. Outputs include quantitative summaries and graphical visualizations such as stacked bar plots, mutation pattern charts, and allele-specific mutation frequency plots (**Fig. 1b and Fig. S1**). For dual-guide systems such as PE3, TwinPE^9^, and PASTE^10^, CRISPRLungo detects and visualizes mutations at both gRNA target sites using a two-site allele plot. Additionally, when a specific intended mutation is specified, CRISPRLungo can classify reads as either precise or imprecise edits.

**Figure 1.**
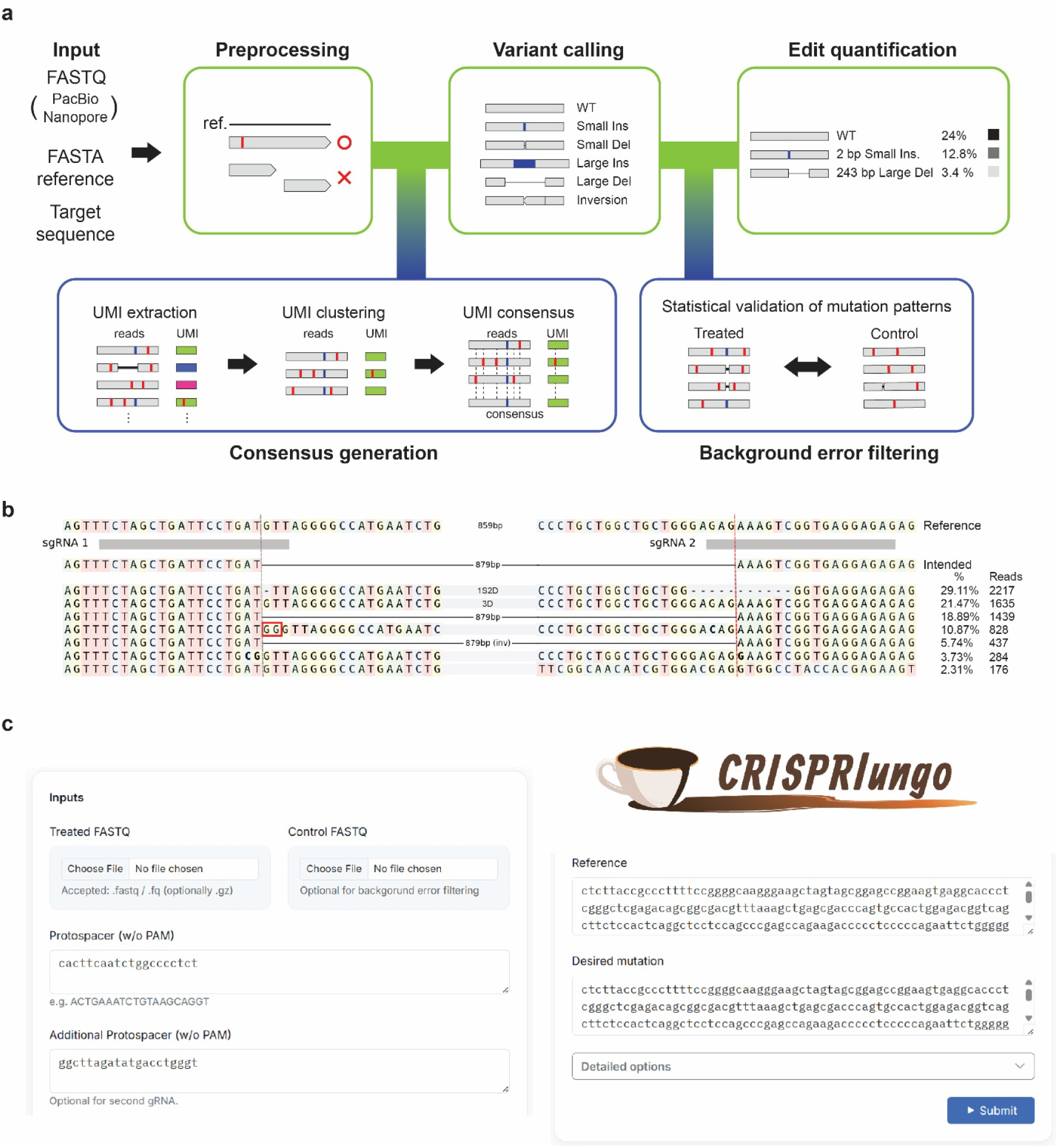
CRISPRLungo: a comprehensive pipeline for CRISPR mutation analysis from long-read sequencing. **a**. Schematic of the CRISPRLungo analysis pipeline. The workflow includes five main stages: preprocessing, variant calling, optional consensus generation, optional background error filtering, and edit quantification with visualization. The modular design allows users to enable UMI-based consensus generation and background error filtering as needed to enhance the accuracy of mutation detection. Vertical red lines indicate sequencing errors, and vertical blue lines indicate insertions. **b**. Representative allele plot produced by CRISPRLungo. Each row denotes an individual allele annotated with mutation type and frequency. Red boxes indicate insertions; short black dashes on a gray background represent small deletions; long horizontal lines depict large deletions. Cleavage sites are shown as vertical dotted lines (gray and red). Mutation types and corresponding read percentages are displayed on the right. **c**. CRISPRLungo web interface. The user-friendly web platform supports the same inputs as the command-line version, allowing direct analysis in the browser without file upload. The platform generates outputs identical to those of the standalone version using the CRISPRLungo.

To encourage broad access, we developed a browser-based version of CRISPRLungo (**Fig. 1c**). A key innovation of our web tool is that all computational analyses occur entirely within the user’s browser, no data are ever uploaded to external servers, ensuring privacy and security for sensitive genomic data. We achieved this through recent WebAssembly technologies, which enable complex bioinformatics tools like our pipeline to run directly in web browsers with near-native performance. Thanks to this unique design, CRISPRLungo requires no software installation, making advanced long-read analysis accessible to researchers regardless of computational resources or expertise. Additionally, Web Workers enable parallel processing, ensuring responsive performance during intensive analyses (**Fig. S2**). Users simply select treated and control reads files (.FASTQ), input the reference sequence(s), protospacer sequence(s), and expected editing sequence, then initiate analysis with a single click. The pipeline automatically filters background errors and generates identical plots and mutation summaries as the command-line version.

### Optimization of CRISPRLungo pipeline

To ensure CRISPRLungo achieves optimal performance across diverse experimental conditions, we systematically evaluated and refined four critical components of the pipeline: (1) validation using simulated datasets with known ground truth, (2) alignment parameter optimization for detecting large structural variants, (3) UMI clustering refinement to minimize mis-clustering errors, and (4) development of statistical tests to filter out false positive events due to sequencing error. Each optimization step was guided by rigorous benchmarking to maximize both sensitivity and specificity.

First, we developed a comprehensive simulation framework to validate CRISPRLungo’s performance by generating long-read sequencing datasets with defined CRISPR-induced edits and platform-specific error profiles (**Fig. 2a**). We synthesized reads modeling small and large indels as well as inversions at predefined cleavage sites and appended adapter sequences with unique molecular identifiers (UMIs). To replicate the non-uniform amplification effects of PCR, we sampled subsets of reads to ensure an average depth of at least 20x. Sequencing errors characteristic of the Nanopore and PacBio platforms were simulated using Badread^28^. Each read was tagged with metadata encoding the underlying mutation, enabling direct comparison between simulated truth and CRISPRLungo analysis outputs (**Fig. S3**). A 5.8 kb fragment from the *BCL11A* gene served as the reference for simulation data.

**Figure 2.**
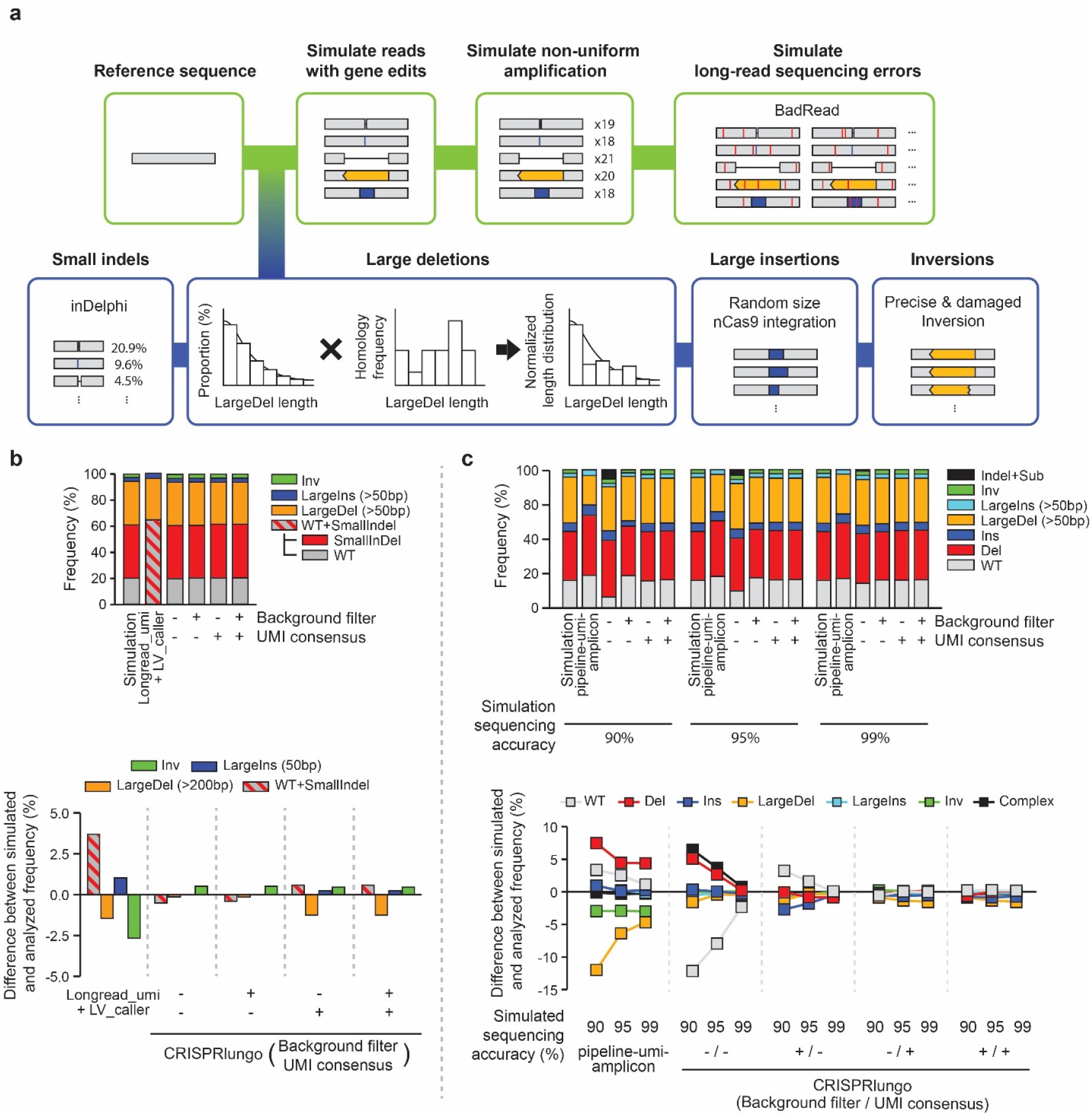
CRISPRLungo accurately quantifies simulated long-read sequencing data. **a**. Scheme of the CRISPR and long-read sequencing simulation pipeline. The simulation process includes CRISPR-induced mutations (small mutations, large deletions, large insertions, and inversions), non-uniform amplification, and sequencing errors. Mutation types are color-coded: black horizontal lines (deletions), orange boxes (inversions), blue boxes (insertions), blue vertical lines (CRISPR-induced substitutions), and red vertical lines (sequencing errors). **b**. Plots showing the frequency of each mutation type detected in PacBio (HiFi) simulation data using CRISPRLungo compared against previously published pipeline. The upper stacked bar plot shows the frequency of each mutation type. The *longread_umi* + *LV_caller* reports WT and SmallIndel as a combined category. The lower plot shows the difference between simulated and analyzed mutation frequencies. The values were calculated by subtracting the simulated frequency from the analyzed frequency. **c**. Plots showing the frequency of each mutation type detected in Nanopore simulation data at three sequencing accuracy levels (90%, 95%, and 99%) using CRISPRLungo compared against previously published pipeline. The upper stacked bar plot and the lower plot follow the same format as in **b**.

Second, we developed an optimized alignment strategy by extending Minimap2. Minimap2, a widely adopted tool for long-read sequencing alignment that is already translated into WebAssembly through the Biowasm project, was selected as the alignment tool for CRISPRLungo to enable seamless integration into the browser-based environment. However, this aligner is suboptimal using default parameters for the analysis of genome editing data. For example, when detecting large deletions, reads flanking the deleted region often produce distant alignments, resulting in split alignments (primary and supplementary). Similarly, inversions disrupt contiguity and lead to complex multi-part alignments. CRISPRLungo was designed to incorporate such supplementary alignments to detect large structural variants. To assess detection fidelity, we simulated deletions and inversions ranging from 100 to 5,000 bp at 100 bp intervals at 98% sequencing accuracy. We observed reduced accuracy for deletions between 2.4-4.0 kb and inversions >3.8 kb (**Fig. S5a**). Asymmetric alignment gaps in cases of large deletions often led to missing supplementary alignments. To address this, we compared the default long-join bandwidth value (20,000) with a reduced value (1,800) tailored to the reference region. The lower value improved alignment recovery and helped mitigate the underestimation of large deletions. However, supplementary alignments for large inversions remained incomplete. To resolve this, we implemented a multi-pass alignment strategy that identifies clipped sequences from primary alignments and re-aligns them independently using Minimap2 (**Fig. S4**). This two-pass method recovered previously missed inversion segments and significantly improved coverage (**Fig. S5b**).

Third, we refined the UMI clustering approach to minimize mis-clustering from sequence similarity. The *pipeline-umi-amplicon* released by Oxford Nanopore employs VSEARCH^28^ for UMI clustering, but its default settings are susceptible to mis-clustering due to loose identity thresholds^29^. To enhance precision, we developed a two-step clustering protocol using VSEARCH, initially grouping by raw UMI sequence, then re-clustering based on the consensus UMI. Results from both rounds were merged to generate the final UMI clusters. We benchmarked multiple identity thresholds on datasets simulated at 90%, 95%, and 99% sequencing accuracy. For each condition, we evaluated mis-clustering by comparing the resulting UMI clusters with known ground-truth assignments and counted the total number of clusters generated. When both clustering steps used a 90% identity threshold, we observed 9 mis-clustered groups. When using 90% for the first round and 95% for the second, mis-clustered groups decreased to 8. The total number of clusters in the second step was 5,626 for the 90%/90% configuration and 5,505 for the 90%/95% configuration. Because the 90/95 setting yielded slightly better accuracy with comparable cluster numbers, we adopted it as the CRISPRLungo default (**Fig. S6**).

Fourth, we developed a two-step statistical procedure to identify true editing patterns despite background noise. The first step evaluates allele count distributions between control and treated data using either chi-squared or Fisher’s exact tests (depending on expected counts), followed by Benjamini-Hochberg (BH) FDR correction to adjust p-values, with a nominal FDR threshold of 0.05 **(Fig. S7A**). While this procedure effectively identifies small mutations, large deletions and insertions are rarely observed in the editing window of control datasets, leading to a lack of power to detect differential editing. To address this limitation, we implemented a secondary statistical test that evaluates the significance of the indel lengths. Specifically, we fitted a shifted negative binomial model to the control datasets and established that indels longer than 10 nt represent a conservative threshold for classifying significant events (**Fig. S7B**). This 10 nt cutoff was validated empirically, as the probability of observing indels exceeding this length in real control samples was extremely low (p < 0.002 across multiple datasets), with only 1 of 158 control alleles exceeding this threshold. These systematic optimizations collectively ensure that CRISPRLungo achieves accurate and reliable mutation detection across the full spectrum of CRISPR editing outcomes.

### Validation of CRISPRLungo

We systematically validated CRISPRLungo’s performance through benchmarking against simulated datasets with known mutation frequencies and comparison to existing quantification approaches across both PacBio and Nanopore platforms. CRISPRLungo operates in four distinct modes depending on whether UMI-based consensus generation and/or background error filtering are applied. To evaluate each mode’s performance, we generated simulated datasets with predefined CRISPR-induced mutation frequencies. We then compared CRISPRLungo results to both the simulated ground truth and outputs from two alternative approaches: (i) *pipeline-umi-amplicon*, a general-purpose tool from Oxford Nanopore to analyze UMI Nanopore sequencing data but not optimized for genome editing analysis, and (ii) the *longread_umi + LV_caller pipeline*, which was proposed in a previous study for CRISPR outcome analysis based long reads. We evaluated *longread_umi* + *LV_caller* on PacBio data and *pipeline-umi-amplicon* on Nanopore data, following the platform-specific implementations described in their original publications. For all comparisons, we simulated datasets with errors corresponding to PacBio HiFi sequencing and separately Nanopore at 90%, 95%, and 99% sequencing accuracy. We assessed accuracy by comparing the ground truth frequencies of each simulated mutation type to the frequencies recovered by each analysis pipeline. Because many studies define large structural variations using a 50-bp threshold and this facilitates comparison with existing approaches, we adopted this cutoff to classify large deletions and insertions^30^.

We first tested CRISPRLungo using simulated PacBio HiFi reads, a scenario with modest sequencing errors, to establish baseline performance (**Fig. 2b**). A critical limitation of the *longread_umi* + *LV_caller* pipeline became immediately apparent: it cannot distinguish between wild-type (WT) reads and small indels, instead grouping them together as a single category (“Small Indel + WT”). This lack of resolution prevents researchers from accurately quantifying editing efficiency, as unedited and edited reads are conflated. By contrast, CRISPRLungo provides precise categorization of WT and small indel reads as separate groups, enabling accurate determination of editing rates. We annotated results accordingly for direct comparison. Consistent with PacBio HiFi’s high intrinsic accuracy (>99.9%)^31^, all four CRISPRLungo modes produced results within 2% of the expected frequency for each mutation class (**Fig. 2b**). In contrast, the *longread_umi* + *LV_caller* pipeline overestimated the “Small Indel + WT” category by 3.8% and failed to detect inversion events.

To assess performance under more challenging conditions, we next evaluated CRISPRLungo on lower-accuracy Nanopore reads (90–99%) where error correction is critical^31^. In this setting, CRISPRLungo’s background error filtering notably improved the agreement between the observed and simulated mutation frequencies (**Fig. 2c**). While UMI consensus generation alone provided substantial error correction, background filtering proved essential for experiments without UMI incorporation, a common scenario in many CRISPR editing studies due to cost or technical constraints. Without UMIs, background filtering reduced false-positive mutation calls by up to 18.7% compared to unfiltered data.

When both the consensus generation and background error filtering options were enabled, the output closely matched the simulated frequencies across all mutation types. By contrast, *pipeline-umi-amplicon* provides only UMI consensus generation and does not perform mutation analysis. Therefore, we applied CRISPRLungo’s mutation calling module to its output. Even with this hybrid approach, *pipeline-umi-amplicon* failed to detect inversion mutations. Collectively, these comprehensive validations demonstrate that CRISPRLungo outperforms existing approaches in both accuracy and mutation coverage, particularly for complex events such as inversions, and provides consistent frequency estimates that closely reflect ground truth in both PacBio and Nanopore simulation datasets. Having established CRISPRLungo’s superior performance on simulated data, we next evaluated its ability to analyze real experimental datasets.

### CRISPRLungo reveals previously undetected mutations in published CRISPR datasets

To demonstrate CRISPRLungo’s ability to extract new biological insights from existing data, we reanalyzed two published long-read sequencing datasets, revealing previously undetected mutation types and validating our quantification against orthogonal methods.

First, we identified previously undiscovered inversion events in a therapeutic genome editing study. We reanalyzed UMI PacBio sequencing datasets from a previously published study^24^, in which SpCas9 was used to edit HSPCs derived from two patients with sickle cell disease (SCD), targeting *BCL11A* and the promoter region of *HBG1/HBG2* (**Fig. 3a**). We applied CRISPRLungo with UMI consensus generation and compared the results with those obtained using the *longread_umi* + *LV_caller* pipeline. For wild-type and small indels grouped together, CRISPRLungo reported a frequency of 80.21 ±□6.72%, while the *longread_umi* + *LV_caller* pipeline reported 78.8□±□4.90% (**Fig. 3b**). For large mutations (>50 bp), including large deletions, large insertions and large complex indels, CRISPRLungo detected 19.64□±□6.74%, compared to 21.23□±□4.89% by the *longread_umi* + *LV_caller* pipeline. The two methods showed strong correlation (r=0.979, p□=□2.15x10^-5^) for large mutation quantification, confirming comparable performance for these mutation types (**Fig. 3c**). Critically, CRISPRLungo detected inversion events at a frequency of 0.15 ± 0.17%, mutations that were completely missed by the original analysis (**Fig. 3d**). Importantly, we confirmed the presence of these inversion-containing reads in the original FASTQ file, validating CRISPRLungo’s detection (**Fig. S8**).

**Figure 3.**
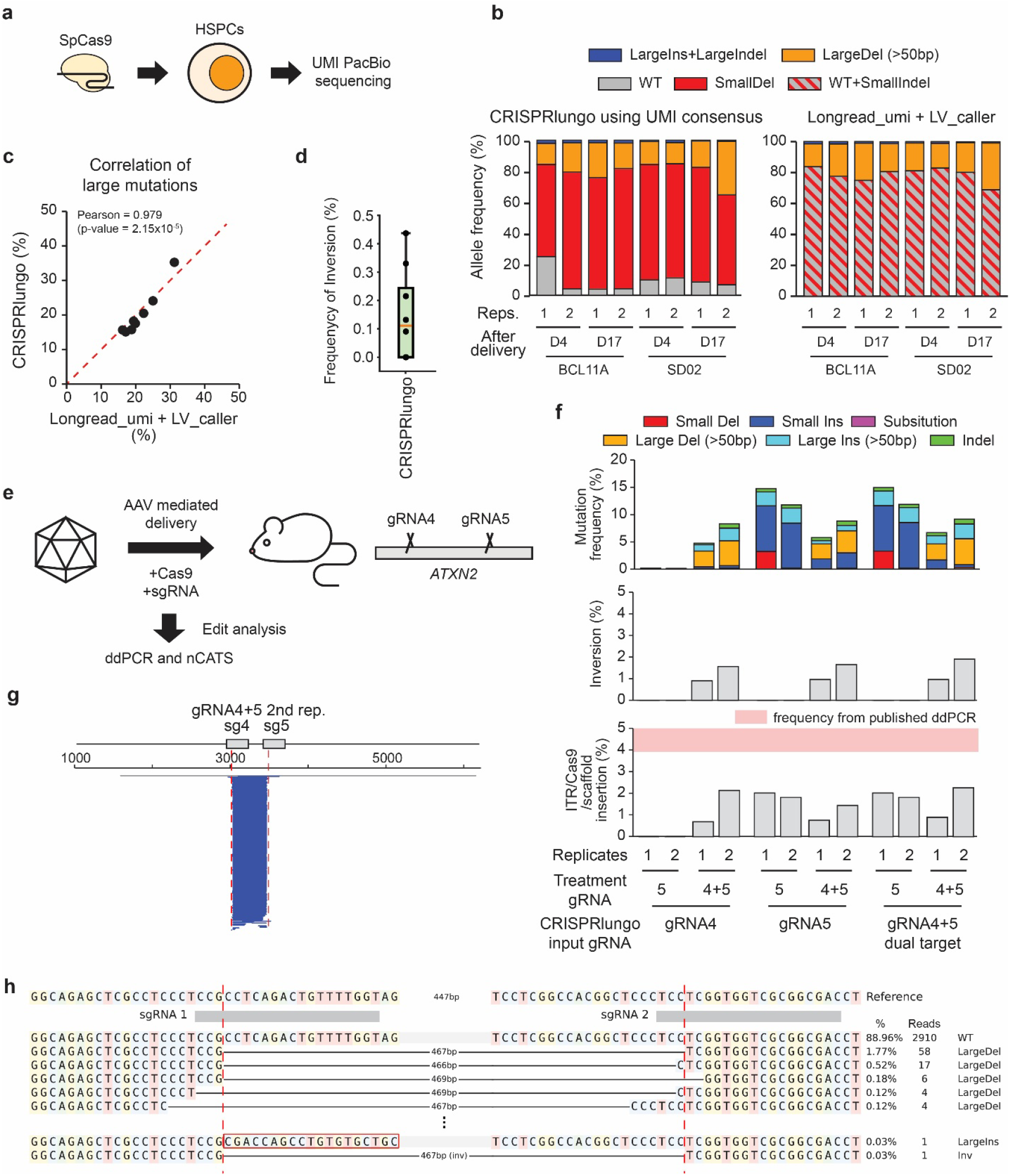
CRISPRLungo re-analysis of published datasets reveals previously undetected gene edits. **a**. Schematic of experimental design using UMI PacBio sequencing on HSPCs edited with SpCas9. **b**. Stacked bar plots comparing mutation frequencies from published data analyzed using CRISPRLungo (with UMI consensus) and the *longread_umi* + *LV_caller* pipeline. **c**. Scatter plot showing the correlation of large deletions (>50 bp) and large insertions between CRISPRLungo and the *longread_umi* + *LV_caller* pipeline (Pearson r = 0.979, p = 2.15x10^-5^). **d**. Box plot showing the frequency of inversions detected by CRISPRLungo (n = 8); these mutations were not detected by the original analysis. **e**. Schematic of the experiment using AAV-mediated delivery of Cas9 and gRNAs targeting *ATXN2*, followed by mutation analysis with ddPCR and nCATS. **f**. Mutation analysis results from nCATS data processed with CRISPRLungo. Top: stacked bar plots showing mutation types and frequencies. Middle: bar graph for inversion frequencies. Bottom: frequency of ITR/Cas9 scaffold integration with reference to published ddPCR data (shaded pink). **g**. Large deletion plot for a dual-gRNA-treated sample (gRNA4+5). Blue bars indicate deletions spanning both cut sites, gray bars indicate bidirectional deletions, and red dotted lines indicate cleavage sites. **h**. Representative allele plot for the gRNA4+5 dual targeting condition. Mutation types are shown along with corresponding read frequencies. Red boxes: insertions; short black lines on gray: small deletions; long lines: large deletions; dotted lines: cleavage sites.

Next, we validated CRISPRLungo’s quantification accuracy using published nCATS (nanopore Cas9⍰targeted sequencing) data from AAV-mediated delivery of Cas9 for in vivo genome editing^32^. In this study, two CRISPR editing strategies targeted the *ATXN2* gene in SCA2 mouse brain: (1) a single guide RNA (gRNA5) designed to induce indels downstream of the CAG repeat expansion, and (2) a dual guide RNA approach (gRNA4+5) designed to delete the entire CAG repeat region. This dataset provided an ideal validation opportunity as editing efficiency and AAV integration events were independently quantified using droplet digital PCR (ddPCR), a gold-standard orthogonal method.

Using CRISPRLungo’s background error filtering capability, we analyzed samples with gRNA5 alone and samples treated with both gRNA4 and gRNA5. For single gRNA5-treated samples, CRISPRLungo reported mutation frequencies of 0.14 ± 0.06% (n=2) at the gRNA4 site, and 13.27 ± 2.11% (n=2) at the gRNA5 site (**Fig. 3f**). For dual-gRNA-treated samples, mutation frequencies were 7.76 ± 2.99% (n=2) for gRNA4, and 8.60 ± 2.62% (n=2) for gRNA5. When analyzing both gRNAs together, CRISPRLungo reported overall mutation frequencies of 9.34 ± 2.40% (n=2). Large deletions were detected in dual-gRNA samples at 3.87 ± 1.28% (n=2), with most showing precise deletion between the two cleavage sites (**Fig. 3g and h**).

To assess AAV integration quantification accuracy, we compared CRISPRLungo with the original ddPCR data. CRISPRLungo identified ITR/Cas9/scaffold insertion events at 1.58 ± 0.97% (n=2) in dual-gRNA samples, lower than the ddPCR-measured integration rates (4-5.5%). While conservative, the detection of these complex insertions with a mean length of 4342 ± 3723 bp demonstrates CRISPRLungo’s capability to identify large structural variants in challenging PCR-free sequencing approaches. This underestimation likely reflects the known length bias of Nanopore sequencing, which underrepresents longer inserts.

Taken together, these reanalyses of both PacBio and Nanopore datasets demonstrate that CRISPRLungo not only provides accurate quantification validated by orthogonal methods but also reveals previously undetected mutation types such as inversions, establishing it as a valuable tool for extensive characterization of CRISPR editing outcomes across diverse experimental contexts.

### CRISPRLungo enables allele-specific gene editing outcome analysis

Accurately assessing gene editing efficiency in patient cells may be challenging when heterozygous mutations and pseudogenes with high sequence homology to the target gene are present. A representative example occurs in patients with Shwachman-Diamond syndrome (SDS). This disorder is caused by biallelic (either homozygous or compound heterozygous) mutations of the *SBDS* gene, which shares high sequence similarity with the pseudogene *SBDSP1*, complicating allele-specific editing analysis. To address this challenge, we developed an additional module for CRISPRLungo to enable read classification based on allele-specific sequence features.

To demonstrate this capability, we analyzed gene editing outcomes in HSPCs from an SDS patient carrying compound heterozygous *SBDS* mutations, following Cas9 editing targeting one mutant allele (c.258+2T>C, *SBDS* allele 1) (**Fig. 4a**). A 2,134 bp fragment was amplified using *SBDS*-specific primers designed to exclude *SBDSP1*, followed by Nanopore sequencing and CRISPRLungo analysis (**Fig. 4b**). In Cas9-treated samples, the overall indel frequency was 47.81%, with 42.85% of reads showing c.258+2T and 9.08% showing c.258+2T>C. By contrast, the mock-treated control exhibited 0.0% indels, with 38.71% c.258+2T, and 61.29% c.258+2T>C (**Fig. 4c and d**). While these results suggest successful editing of *SBDS* allele 1, the minimal sequence differences between the two *SBDS* alleles and potential pseudogene contamination prevented definitive assessment of off-target editing at *SBDS* allele 2. Indeed, IGV^33^ visualization revealed reads with *SBDSP1*-specific variants, confirming pseudogene amplification despite gene-specific primers (**Fig. S9**).

**Figure 4.**
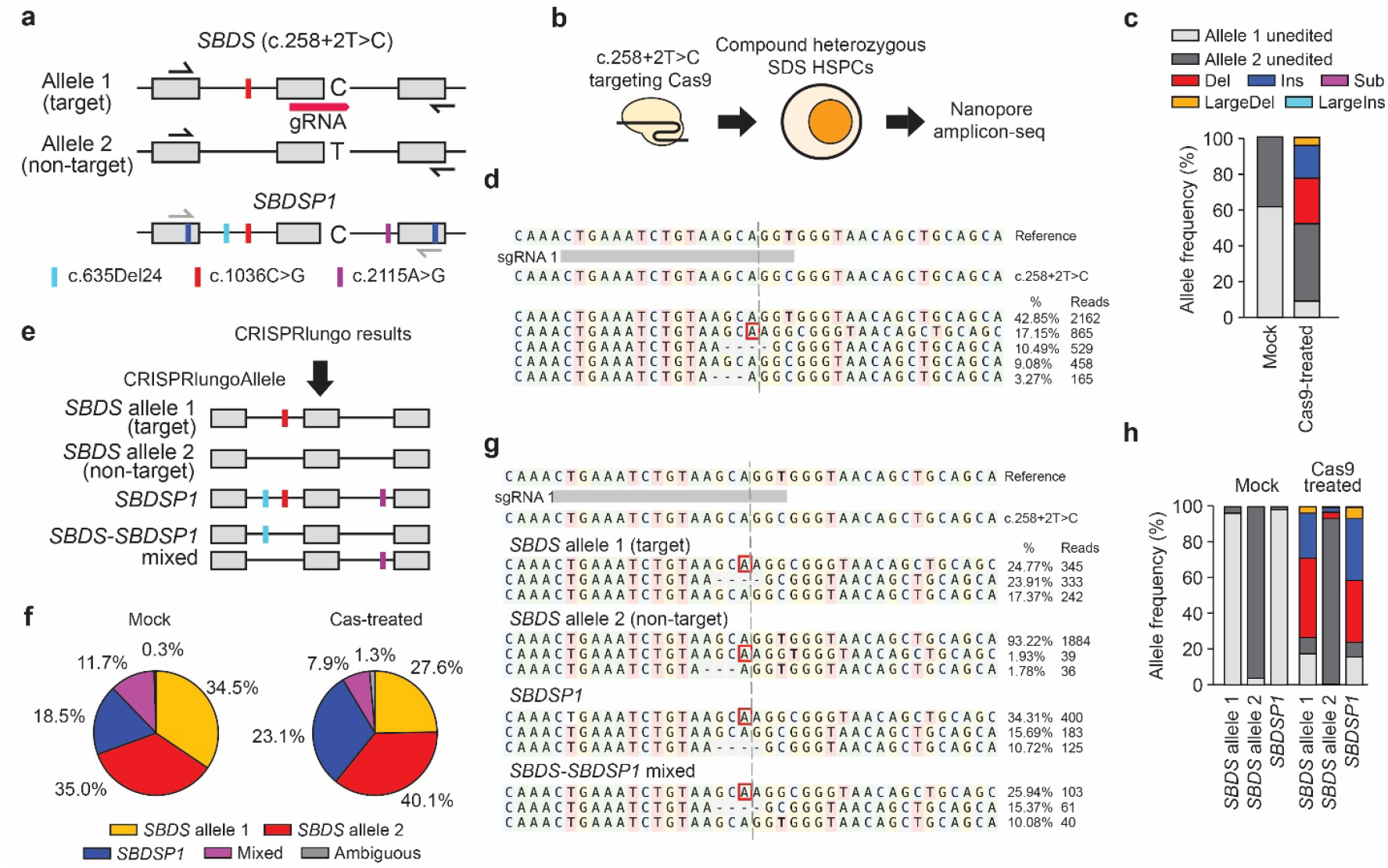
CRISPRLungo allele-specific analysis of gene editing in *SBDS* compound heterozygous patient cells with *SBDSP1* pseudogene read contamination. **a**. Schematic representation of *SBDS* and *SBDSP1* alleles in SDS patient cells. Black half arrows represent primers, gray arrows indicate primer mismatches, and colored bars indicate distinguishing sequence variants: sky blue (c.635Del24), red (c.1036C>G), and purple (c.2115A>G). The navy blue bars mark the pseudogene specific mutations, and the red horizontal bar represents the gRNA. **b**. Schematic of the experiment delivering SpCas9 to HSPCs from SDS patients, followed by Nanopore amplicon sequencing. **c**. Stacked bar plot of mutation frequencies in mock and Cas9-treated samples. **d**. Representative allele plot for Cas9-treated samples. Red boxes indicate insertions; short black dashes on a gray background represent small deletions; Cleavage sites are shown as vertical dotted lines (gray). **e**. Schematic of CRISPRLungo allele classification for *SBDS* allele 1 (nothing), *SBDS* allele 2 (red), *SBDSP1* (sky blue, red, and purple), and *SBDS–SBDSP1* mixed (only sky blue or only purple). **f**. Pie charts showing the distribution of allele groups in mock and Cas9-treated samples. **g**. Representative allele plots of each classified group. **h**. Stacked bar plots of mutation frequencies for each allele group.

To improve specificity and achieve true allele-specific resolution, we developed a custom classification strategy exploiting sequence differences between the gene and pseudogene. We leveraged three distinguishing sequence variants, c.635del24, c.1036C>G, and c.2115A>G, which are differently present in *SBDS* and *SBDSP1* (**Fig. 4a and e**). We implemented this strategy as an integrated module within CRISPRLungo, which we termed CRISPRLungoAllele. This module processes the aligned reads from the main pipeline and applies custom variant-based classification rules to segregate reads by their allelic origin. Using CRISPRLungoAllele, we classified reads into five categories: (i) *SBDS* allele 1 (c.1036C>G only), (ii) *SBDS* allele 2 (none of the three variants), (iii) *SBDSP1* allele (all three variants), (iv) *SBDS*–*SBDSP1* mixed alleles, containing *SBDSP1*-specific variants on one end and WT *SBDS* sequence on the other, and (v) ambiguous alleles that do not belong to any specific category.

In the mock sample, the distribution of classified reads was as follows: *SBDS* allele 1, 34.5%; *SBDS* allele 2, 35.0%; *SBDSP1*, 18.5%; and mixed, 11.7% (**Fig. 4f**). Cas9 treatment shifted these proportions to 27.6%, 40.1%, 23.1%, and 7.8%, respectively. These results indicate that despite the use of *SBDS*-specific primers, amplification of *SBDSP1* and generation of chimeric reads could not be fully avoided. Indel analysis revealed minimal off-target editing at *SBDS* allele 2 (0.0% in mock, 6.53% in treated), suggesting limited editing activity on the untargeted allele. In contrast, *SBDS* allele 1 showed a dramatic indel increase from 0.0 % to 73.4%, confirming efficient allele-preferential editing. Reads classified as *SBDSP1* also showed elevated indel rates (0.0% to 76.2%), reflecting identical sequence at the Cas9 target site between *SBDS* and *SBDSP1* (**Fig. 4g, h, and Fig. S10**).

These findings establish that CRISPRLungo enables high-resolution, allele-specific analysis of gene editing in complex heterozygous patient samples, even in the presence of pseudogenes and chimeric amplification artifacts. This capability provides critical distinction between intended on-target editing and off-target effects, essential for evaluating the precision of therapeutic genome editing strategies.

## DISCUSSION

The rapid evolution of CRISPR-based genome editing technologies has broadened the landscape of possible allelic outcomes, necessitating more comprehensive analytical approaches. Techniques such as PE3 and TwinPE, which induce editing with multiple gRNAs with distinct nick positions, have further increased analytical complexity. In parallel, growing attention to unintended structural variant outcomes from genome editing has underscored the need for more versatile tools. While long-read sequencing platforms have become widely adopted, no existing analysis pipeline provides comprehensive support for all relevant CRISPR editing contexts.

In this study, we introduced CRISPRLungo, a long-read sequencing analysis pipeline designed to accommodate an extensive spectrum of CRISPR-induced mutations. CRISPRLungo demonstrated high accuracy in simulation benchmarks, detecting a wide range of mutational events with minimal deviation from ground truth. Furthermore, it successfully analyzed results from PCR-free methodologies such as nCATS, demonstrating compatibility across diverse experimental designs. To support detailed investigations, including allele-specific mutation tracking and identification of chimeric amplification products, CRISPRLungo incorporates an optional module for read-level classification based on user-defined sequence features, including naturally occurring variants for distinguishing alleles and pseudogenes or artificial barcodes for multiplexed experiments, providing essential capabilities for evaluating therapeutic editing outcomes.

Our simulations focused on distinguishing true mutations from sequencing background noise. However, we acknowledge that real experimental conditions may introduce additional biases not explicitly captured in our models. PCR amplification produces sequence- and length-dependent biases, and long-read sequencing exhibits read length-associated artifacts^23^. These variables were not incorporated into our benchmarking but may influence editing outcome interpretation depending on library construction methods. UMI-based long-read sequencing and optimized PCR protocols represent promising approaches for mitigating such biases in future applications.

Technical challenges were also evident in our analysis of patient-derived *SBDS* samples, where 7–12% of reads were classified as *SBDS*–*SBDSP1* mixed alleles. Two mechanisms could explain these chimeric sequences. First, genomic rearrangements such as inversion events between *SBDS* and its pseudogene *SBDSP1*, both located on chromosome 7 in opposing orientations, could result in fusion junctions^34^. Second, PCR-mediated chimeras, formed by incomplete extension and template switching, likely account for the majority of these reads^35^. The presence of *SBDS*–*SBDSP1* mixed reads even in mock-treated controls strongly suggests a technical rather than biological origin. Additional evidence of intra-*SBDS* chimeras was found in reads annotated as *SBDS* allele 1 and allele 2, which frequently carried sequence markers from the alternate allele. This likely arises from primer designs where discriminating variants for *SBDS* and *SBDSP1* are present at both ends, allowing precise identification of inter-locus chimeras, while *SBDS*-specific alleles are distinguished by markers on only one end. Future improvements to primer design, incorporating allele-discriminating bases at both termini, would further enhance specificity in allele-resolved editing quantification.

Beyond analytical precision, user access was a key consideration in CRISPRLungo’s development. We implemented a browser-based version that executes entirely on the client-side. This design eliminates complex installations and, more importantly prevents data uploads, ensuring data privacy. The webtool enables users to conduct end-to-end analyses without requiring computational expertise. Using Nanopore sequencing and the CRISPRLungo web tool, users can analyze DNA sequences from samples within hours. Due to computational limitations of browser environments, resource-intensive tasks such as UMI-based consensus generation are currently not supported in the web tool. However, with the emergence of the WebGPU API, it is now feasible to implement GPU-accelerated workflows within the browser. Integration of UMI consensus generation and genome-wide alignment into the web platform represents our next development priority. Importantly, CRISPRLungo eliminates the need for individual laboratories to develop custom analysis pipelines, saving substantial time and resources while improving reproducibility across the field.

In conclusion, CRISPRLungo provides a unified platform for analyzing long-read sequencing data from CRISPR-edited cells. The pipeline delivers highly accurate mutation detection, supports allele-level resolution, and identifies a broader array of editing outcomes than existing tools (**Supplementary Note 1**). We expect CRISPRLungo to establish a new standard for long-read sequencing analysis in CRISPR genome editing, supporting both experimental research and therapeutic development.

## METHODS

### CRISPRLungo analysis pipeline

The CRISPRLungo pipeline was implemented in Python3 (v3.10) and consists of modular components for alignment, UMI processing, statistical mutation analysis and visualization. Currently CRISPRLungo analyzes one control and one treated sample at a time. When multiple replicates exist, we recommend combining them before analysis. This approach is justified under the assumptions of the statistical tests used in the pipeline: Fisher exact and Chi-squared test.

1. <u>Alignment module</u>: CRISPRLungo’s alignment module utilizes Minimap2 (v2.22)^1^ with parameters optimized for long-read CRISPR analyses: “-ax map-ont -p 0.5 -r {0.3 × reference_length}, {500 (or 0.3 × reference_length if the latter was < 500)}” option. To capture large structural variants, we implemented a multi-pass alignment strategy: primary alignments are processed first, then clipped segments ≥100 bp are iteratively realigned until no additional clipped sequences remain. All resulting SAM files are merged using the CRISPRLungo_minimap.py script to create a comprehensive alignment profile.
2. <u>UMI consensus generation</u>: When UMI-based error correction is enabled (--umi), CRISPRLungo extracts UMI sequences from positions annotated in the input FASTA reference file, such as: “AAGCAGAAGACGGCATACGTGAT(NNNYRNNNYRNNNYRNNN)TCCAA…atggcaac(NNNYRNNNYRNNNYRNNN)GATCTCGGTGGTCGCCGTATCATT”. After aligning FASTQ reads, bases aligned at those positions are extracted and assigned as each read’s UMI sequence. We implemented a two-step clustering approach using VSEARCH (v2.26.0)^2^ to minimize mis-clustering errors. First using “--minseqlength {umi_len - 1} -- maxseqlength {umi_len + 1} --qmask none --cluster_fast {input_file} --gapopen 0E/5I -- gapext 0E2I --mismatch -8 --match 6 --iddef 0 --minwordmatches 0 -id 0.9”, then repeating with the same options, except for the -id flag that is increased to 0.95. Reads are then grouped according to the second-round clustering results, and consensus sequences are generated using the pyspoa (v0.2.1, https://github.com/nanoporetech/pyspoa) to produce error-corrected reads.
3. <u>Background error filtering</u>: For experiments with control samples (--control), CRISPRLungo implements statistical filtering to distinguish true mutations from technical artifacts. The pipeline extracts mutation patterns from both treated and control samples within the user-specified cleavage window (--window). Mutations not observed in control samples exceeding 10nt in length are considered significant mutations. To establish this cutoff, insertion and deletion length distributions from multiple untreated control datasets (UMI PacBio *BCL11A* and nCATS) were modeled using shifted negative binomial models. The estimated probability of observing indels longer than 10 bp in control samples was extremely low (p = 8 × 10^−^□ to 2 × 10^−3^ across datasets), with only 1 of 158 control alleles exceeding this threshold. Thus, >10 bp was adopted as a conservative criterion for classifying significant indels. For remaining mutations in the edited sample, a 2x2 contingency table was constructed representing the presence or absence of the mutations in edited and control samples, and p-values are calculated using either a chi⍰squared test (scipy.stats.chi2_contingency) or a Fisher’s exact test (scipy.stats.fisher_exact) based on whether all expected counts exceed 5^3^. P-values are assigned to each mutation pattern and a Benjamini-Hochberg correction with an FDR of 0.05 is applied to identify the significant mutations. This approach effectively controlled the empirical FDR (0.008 at 90% sequencing accuracy and 0.004 at 99% sequencing accuracy) in simulations. Among all identified mutation patterns, only those deemed significant or matching the user-defined intended reference sequence were included in downstream analysis and visualization. Only mutations spanning the user-specified cleavage window (--window) are analyzed for CRISPR-induced mutations.
4. <u>Visualization</u>: All interactive visualizations are generated using Plotly for web-based interactivity, while static allele plots and tornado plots are created using Matplotlib for publication-quality figures.

### CRISPRLungo web version

The CRISPRLungo web tool is hosted on GitHub Pages and styled with Bootstrap (https://getbootstrap.com/). To ensure non-blocking execution in web browsers, the tool performs time-consuming analyses (alignment and background error filtering process) by instantiating dedicated Web Workers (**Fig. S2**). While the analysis is running, the browser continuously receives updates from the Web Workers and displays the current progress to the user. Upon submission of FASTQ files, the target sequence, the reference sequence, and the intended reference sequence, the tool first validates all inputs. Treated and control FASTQ files are aligned in parallel. For alignment, a dedicated Web Worker is instantiated in the browser and the Biowasm (https://biowasm.com/) minimap2 module is imported. The alignment workflow is identical to the command line CRISPRLungo pipeline. Once alignment is complete, background error filtering is performed. The background error filtering pipeline was rewritten in Rust and compiled to WebAssembly for high-performance in-browser execution. Results are visualized using Plotly.js (https://plotly.com/javascript/) and D3.js (https://d3js.org/). Due to memory limitations of web browsers, UMI-based consensus generation is currently not supported in the browser version. All other functionalities, including statistical filtering and visualization, are fully supported.

### CRISPR-induced mutations and long-read sequencing simulation pipeline

In our simulation pipeline, each mutation pattern was generated as follows: i) Small indels were predicted using inDelphi^4^, and the resulting mutation frequencies were normalized to match the specified small indel frequency. ii) To simulate large deletions, we generated a half-normal distribution based on the defined variance and mean parameters, reflecting the expected large-deletion frequency and length. For each size bin, the probability of microhomology⍰mediated repair was calculated as the fraction of all possible large deletions within that range that have at least 2 bp of microhomology at their breakpoint junctions. Subsequently, the adjusted large-deletion proportion curve was obtained by multiplying the half-normal distribution with the microhomology-based probability across each size range. Based on this adjusted large-deletion proportion curve, random start positions, including the cleavage site, were sampled to simulate large deletions. Reads were then generated to match the specified large-deletion frequency. Because inDelphi does not predict deletions between 20 bp and 200 bp, 20∼200bp deletion reads were additionally generated according to the adjusted large-deletion proportion curve. iii) large insertions were created by inserting Cas9 sequences of random lengths (10–1,000 bp) at the cleavage site. iv) Inversions were simulated from predefined sequence segments and 20% of inversions were truncated at both ends by random lengths to model end-resection. Each read was tagged with a UMI as follows: the Nanopore simulation reads received the UMI sequence TTTVVVVTTVVVVTTVVVVTTVVVVTTT and the PacBio simulation reads received NNNYRNNNYRNNNYRNNN.

Additionally, all reads in both simulation datasets were flanked with adapter sequences: a 5′ extension of: GGTGCTGAAGAAAGTTGTCGGTGTCTTTGTGTTAACCGTATCGTGTAGAGACTGCGTAGGTTT and a 3′ extension of:

AAACACTCGCACTGACTCGATCACTGGTTAACACAAAGACACCGACAACTTTCTTCAGCACC.

After generating these mutation-containing reads, non-uniform amplification was simulated by randomly subsampling to an average depth of 20×. Finally, long⍰read sequencing errors were introduced using Badread^5^. PacBio reads were simulated using “--error_model pacbio2021 -- qscore_model pacbio2021 --identity 30,3” and Nanopore reads were simulated using “--identity 99,99.9,0.5”, “--identity 95,99,2.5”, and “--identity 90,98,5” with the default error model.

### Validation of CRISPRLungo against existing pipelines

The PacBio simulation dataset was analyzed using the *longread_umi* pipeline^6^ and *LV_caller*^7^, with all parameters set to their documented defaults except that the *LV_caller* -rxn flag was set to 2. The Nanopore simulation dataset was analyzed with *pipeline-umi-amplicon*, modifying only the min_reads_per_cluster threshold to 5 to match CRISPRLungo’s setting. CRISPRLungo was run on both simulation datasets using its default options.

### Reanalysis of published long-read sequencing datasets

The published PacBio sequencing dataset was downloaded from PRJNA780655^7^. The PacBio dataset was analyzed using *longread_umi* (https://github.com/SorenKarst/longread_umi) and *LV_caller* (Zenodo, access code 6805011) with default settings except that the *LV_caller* -rxn flag was set to 2 to resolve a version cofliction. The same PacBio dataset was also analyzed using CRISPRLungo, and deletion events were reclassified into small (<50 bp), intermediate (50–200 bp), and large (>200 bp) deletions for direct comparison. The published nCATS dataset was downloaded from PRJNA916868^8^. The nCATS dataset was analyzed using CRISPRLungo, with the Cas9-enriched sequence used as the reference. The CRISPRLungo results were compared to the ddPCR measurements reported in the original manuscript to validate quantification accuracy.

### Cell culture and electroporation

CD34^+^ HSPCs were thawed and cultured in GMP-grade Stem Cell Growth Medium (SCGM) (CellGenix, 20806-0500), supplemented with 100 ng/mL preclinical thrombopoietin (TPO) (CellGenix, 1417-050), 100 ng/mL preclinical stem cell factor (SCF) (CellGenix, 1418-050), and 100 ng/mL preclinical FMS-like tyrosine kinase 3 ligand (FLT3L) (CellGenix, 1415-050). Electroporation was performed 48 hours post-thaw using the Amaxa 4D-Nucleofector system (P3 Primary Cell 4D-Nucleofector® X Kit S, Lonza, V4XP-3032), with Alt-R™ SpCas9 Nuclease V3 (IDT, 1081058) complexed with sgRNA as a ribonucleoprotein (RNP) complex. Genomic DNA was extracted 5 days after electroporation using the DNeasy Blood & Tissue Kit (Qiagen, 69504).

### Library preparation and Nanopore sequencing

Targeted genomic regions were amplified using the following PCR mixture: 40 ng of genomic DNA, 2.5 μL each of 10 μM forward and reverse primers, 25 μL Platinum™ SuperFi™ DNA Polymerase (Thermo Fisher Scientific, 12351010), and nuclease-free water to a final volume of 50 μL. PCR cycling conditions were as follows: 98□°C for 30 seconds; 4 cycles of 98□°C for 10 seconds, 60□°C for 10 seconds, and 72□°C for 2 minutes; followed by a final extension at 72□°C for 5 minutes. PCR products were purified using 0.9× SPRISelect beads (Beckman Coulter, B23319). Nanopore libraries were prepared using the Ligation Sequencing Kit V14 (Oxford Nanopore Technologies, SQK-LSK114) following the manufacturer’s protocol, and sequencing was performed on a MinION device using an R10.4.1 flow cell (Oxford Nanopore Technologies, FLO-MIN114). The fastq files were generated using Dorado (v1.1.0) with super high accuracy model. The resulting Nanopore data were analyzed using CRISPRLungo.

## Data Availability

In this study, all published datasets used were obtained from the Sequence Read Archive (SRA), with accession IDs provided in the manuscript (**Table S1**). The patient-derived CRISPR treated data have also been deposited in the SRA (PRJNA1345895) and will be made publicly available upon publication of this manuscript.

## Code Availability

CRISPRLungo (v0.1) is freely available at https://github.com/pinellolab/CRISPRLungo. The repository includes comprehensive documentation, example datasets, and tutorials for both command-line and web implementations. A fully functional web interface requiring no installation is accessible at https://pinellolab.github.io/CRISPRLungo.

## Author contribution

Conceptualization: G.H., D.E.B., and L.P.; Software development and validation: G.H. and B.V.; Data acquisition: G.H. and B.V.; Investigation: G.H., J.Z., J.M., A.S., and T.B.; Writing – original draft: G.H.; Writing – review and editing: G.H., B.V., T.B., D.E.B., and L.P.; Funding acquisition: D.E.B. and L.P.; Supervision: D.E.B. and L.P.

## Acknowledgements

The authors thank funding from R01HG013618 (L.P. and D.B), 1R35HG010717(L.P. and D.B.), UM1HG012010 (L.P. and,D.B.) Rappaport MGH Research Scholar Award 2024-2029 (L.P.) This work was supported by the Sejong Fellowship through the National Research Foundation of Korea (Grant No. RS-2024-00343250). Finally, we thank members of the Pinello and Bauer lab for providing important feedback on this project.

## REFERENCES

1. Mali, P. et al. RNA-guided human genome engineering via Cas9. Science 339, 823–826 (2013).

2. Jinek, M. et al. A programmable dual-RNA-guided DNA endonuclease in adaptive bacterial immunity. Science 337, 816–821 (2012).

3. Gaudelli, N.M. et al. Programmable base editing of A*T to G*C in genomic DNA without DNA cleavage. Nature 551, 464–471 (2017).

4. Komor, A.C., Kim, Y.B., Packer, M.S., Zuris, J.A. & Liu, D.R. Programmable editing of a target base in genomic DNA without double-stranded DNA cleavage. Nature 533, 420–424 (2016).

5. Ran, F.A. et al. Double nicking by RNA-guided CRISPR Cas9 for enhanced genome editing specificity. Cell 154, 1380–1389 (2013).

6. Yan, J. et al. Improving prime editing with an endogenous small RNA-binding protein. Nature 628, 639–647 (2024).

7. Anzalone, A.V. et al. Search-and-replace genome editing without double-strand breaks or donor DNA. Nature 576, 149–157 (2019).

8. Choi, J. et al. Precise genomic deletions using paired prime editing. Nat Biotechnol 40, 218–226 (2022).

9. Anzalone, A.V. et al. Programmable deletion, replacement, integration and inversion of large DNA sequences with twin prime editing. Nat Biotechnol 40, 731–740 (2022).

10. Yarnall, M.T.N. et al. Drag-and-drop genome insertion of large sequences without double-strand DNA cleavage using CRISPR-directed integrases. Nat Biotechnol 41, 500–512 (2023).

11. Ansori, A.N. et al. Application of CRISPR-Cas9 genome editing technology in various fields: A review. Narra J 3, e184 (2023).

12. Levesque, S. & Bauer, D.E. CRISPR-based therapeutic genome editing for inherited blood disorders. Nat Rev Drug Discov (2025).

13. Zhu, H., Li, C. & Gao, C. Applications of CRISPR-Cas in agriculture and plant biotechnology. Nat Rev Mol Cell Biol 21, 661–677 (2020).

14. Clement, K. et al. CRISPResso2 provides accurate and rapid genome editing sequence analysis. Nat Biotechnol 37, 224–226 (2019).

15. Hunt, J.M.T., Samson, C.A., Rand, A.D. & Sheppard, H.M. Unintended CRISPR-Cas9 editing outcomes: a review of the detection and prevalence of structural variants generated by geneediting in human cells. Hum Genet 142, 705–720 (2023).

16. Fu, Y. et al. High-frequency off-target mutagenesis induced by CRISPR-Cas nucleases in human cells. Nat Biotechnol 31, 822–826 (2013).

17. Kosicki, M., Tomberg, K. & Bradley, A. Repair of double-strand breaks induced by CRISPR-Cas9 leads to large deletions and complex rearrangements. Nat Biotechnol 36, 765–771 (2018).

18. Hwang, G.H. et al. Large DNA deletions occur during DNA repair at 20-fold lower frequency for base editors and prime editors than for Cas9 nucleases. Nat Biomed Eng 9, 79–92 (2025).

19. Giannoukos, G. et al. UDiTaS, a genome editing detection method for indels and genome rearrangements. BMC Genomics 19, 212 (2018).

20. Yin, J. et al. Optimizing genome editing strategy by primer-extension-mediated sequencing. Cell Discov 5, 18 (2019).

21. Wen, W. et al. Effective control of large deletions after double-strand breaks by homology-directed repair and dsODN insertion. Genome Biol 22, 236 (2021).

22. Carvalho, A.B., Kim, B.Y. & Uno, F. Strong sequencing bias in Nanopore and PacBio prevents assembly of Drosophila melanogaster Y-linked genes. bioRxiv (2025).

23. Dabney, J. & Meyer, M. Length and GC-biases during sequencing library amplification: a comparison of various polymerase-buffer systems with ancient and modern DNA sequencing libraries. Biotechniques 52, 87–94 (2012).

24. Park, S.H. et al. Comprehensive analysis and accurate quantification of unintended large gene modifications induced by CRISPR-Cas9 gene editing. Sci Adv 8, eabo7676 (2022).

25. Karst, S.M. et al. High-accuracy long-read amplicon sequences using unique molecular identifiers with Nanopore or PacBio sequencing. Nat Methods 18, 165–169 (2021).

26. Cull, A.H., Kent, D.G. & Warren, A.J. Emerging genetic technologies informing personalized medicine in Shwachman-Diamond syndrome and other inherited BMF disorders. Blood 144, 931–939 (2024).

27. Li, H. Minimap2: pairwise alignment for nucleotide sequences. Bioinformatics 34, 3094–3100 (2018).

28. Rognes, T., Flouri, T., Nichols, B., Quince, C. & Mahe, F. VSEARCH: a versatile open source tool for metagenomics. PeerJ 4, e2584 (2016).

29. Amstler, S. et al. Nanopore sequencing with unique molecular identifiers enables accurate mutation analysis and haplotyping in the complex lipoprotein(a) KIV-2 VNTR. Genome Med 16, 117 (2024).

30. Mahmoud, M. et al. Structural variant calling: the long and the short of it. Genome Biol 20, 246 (2019).

31. Logsdon, G.A., Vollger, M.R. & Eichler, E.E. Long-read human genome sequencing and its applications. Nat Rev Genet 21, 597–614 (2020).

32. Simpson, B.P., Yrigollen, C.M., Izda, A. & Davidson, B.L. Targeted long-read sequencing captures CRISPR editing and AAV integration outcomes in brain. Mol Ther 31, 760–773 (2023).

33. Robinson, J.T., et al. Integrative genomics viewer. Nat Biotechnol 21, 24–26 (2011).

34. Boocock, G.R. et al. Mutations in SBDS are associated with Shwachman-Diamond syndrome. Nat Genet 33, 97–101 (2003).

35. Judo, M.S., Wedel, A.B. & Wilson, C. Stimulation and suppression of PCR-mediated recombination. Nucleic Acids Res 26, 1819–1825 (1998).

## METHODS REFERENCES

1. Li, H. Minimap2: pairwise alignment for nucleotide sequences. Bioinformatics 34, 3094–3100 (2018).

2. Rognes, T., Flouri, T., Nichols, B., Quince, C. & Mahe, F. VSEARCH: a versatile open source tool for metagenomics. PeerJ 4, e2584 (2016).

3. Nowacki, A. Chi-square and Fisher’s exact tests (From the “Biostatistics and Epidemiology Lecture Series, Part 1”). Cleve Clin J Med 84, e20–e25 (2017).

2. Shen, M.W. et al. Predictable and precise template-free CRISPR editing of pathogenic variants. Nature 563, 646–651 (2018).

3. Wick, R.R. Badread: simulation of error-prone long reads. The Journal of Open Source Software 4, 1316 (2019).

4. Karst, S.M. et al. High-accuracy long-read amplicon sequences using unique molecular identifiers with Nanopore or PacBio sequencing. Nat Methods 18, 165–169 (2021).

5. Park, S.H. et al. Comprehensive analysis and accurate quantification of unintended large gene modifications induced by CRISPR-Cas9 gene editing. Sci Adv 8, eabo7676 (2022).

6. Simpson, B.P., Yrigollen, C.M., Izda, A. & Davidson, B.L. Targeted long-read sequencing captures CRISPR editing and AAV integration outcomes in brain. Mol Ther 31, 760–773 (2023).

