## Supplementary material for "Analyzing long-read CRISPR experiments with CRISPRLungo": Supplemetnary Figures and Note

**CRISPRlungo software for long read sequencing genome editing analysis**

**Affiliations:** ^1^Molecular Pathology Unit, Center for Cancer Research, Massachusetts General Hospital, Department of Pathology, Harvard Medical School, Boston, MA, USA ^2^Division of Hematology/Oncology, Boston Children’s Hospital, Department of Pediatric Oncology, Dana-Farber Cancer Institute, Harvard Stem Cell Institute, Department of Pediatrics, Harvard Medical School, Boston, MA, USA ^3^Department of Chemistry, Hanyang University, Seoul, 04763, South Korea

**This PDF file includes:**

Supplementary Figure 1-11, Supplementary Note 1

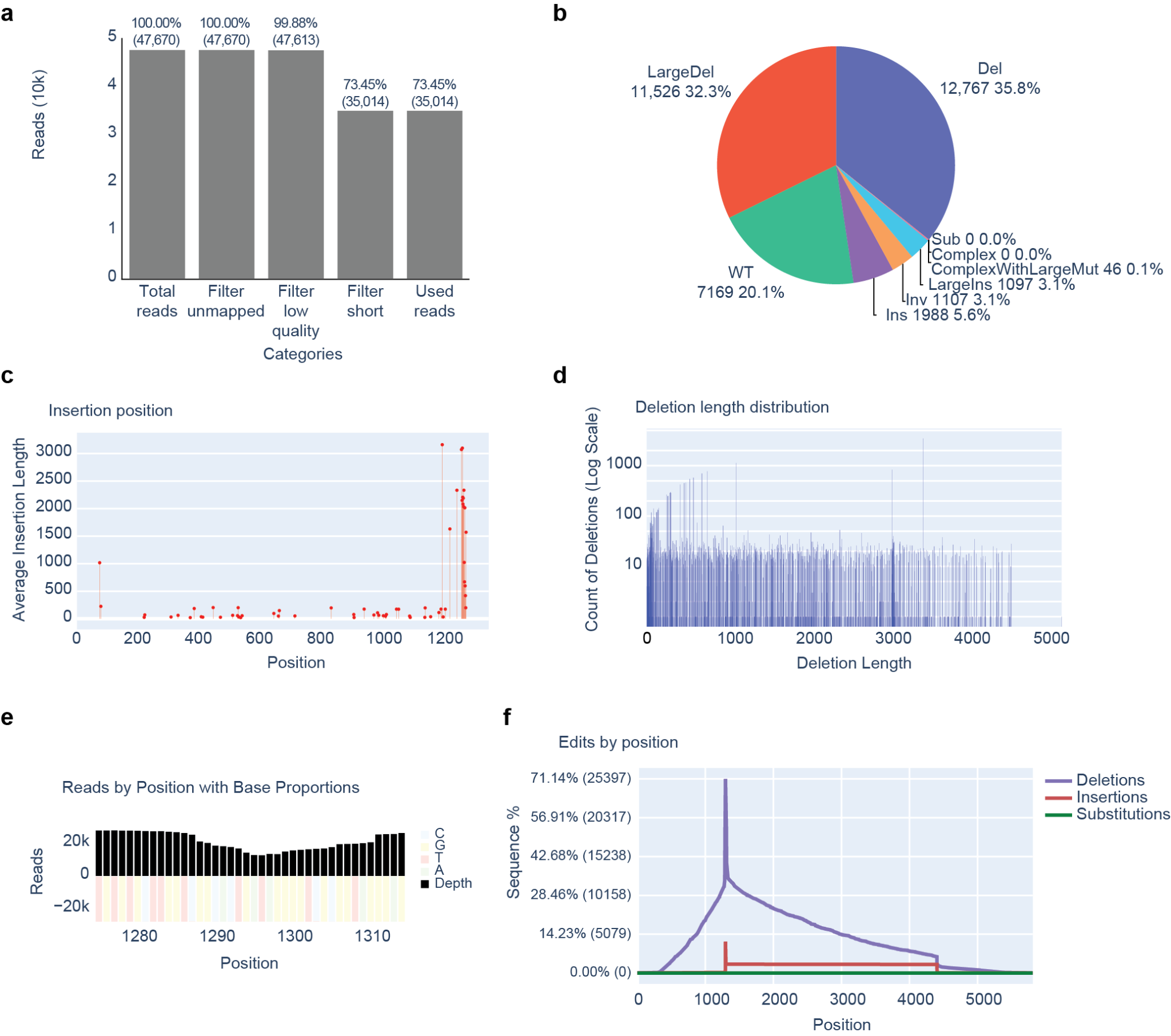

**Supplementary Figure 1. CRISPRlungo output examples.**

**a.** Bar plot of preprocessing summary results. The number of reads passing each filtering step is shown. **b**. Pie chart representing the proportion of each edit type. **c**. Plot of average insertion length at each genomic position. **d**. Distribution plot of deletion counts by deletion length. **e**. Plot of base proportion and read depth across positions within the window. Indels are excluded from the base proportion panel. **f**. Distribution of insertion, deletion, and substitution frequencies across positions. This example was generated using nanopore simulation data with 99% accuracy.

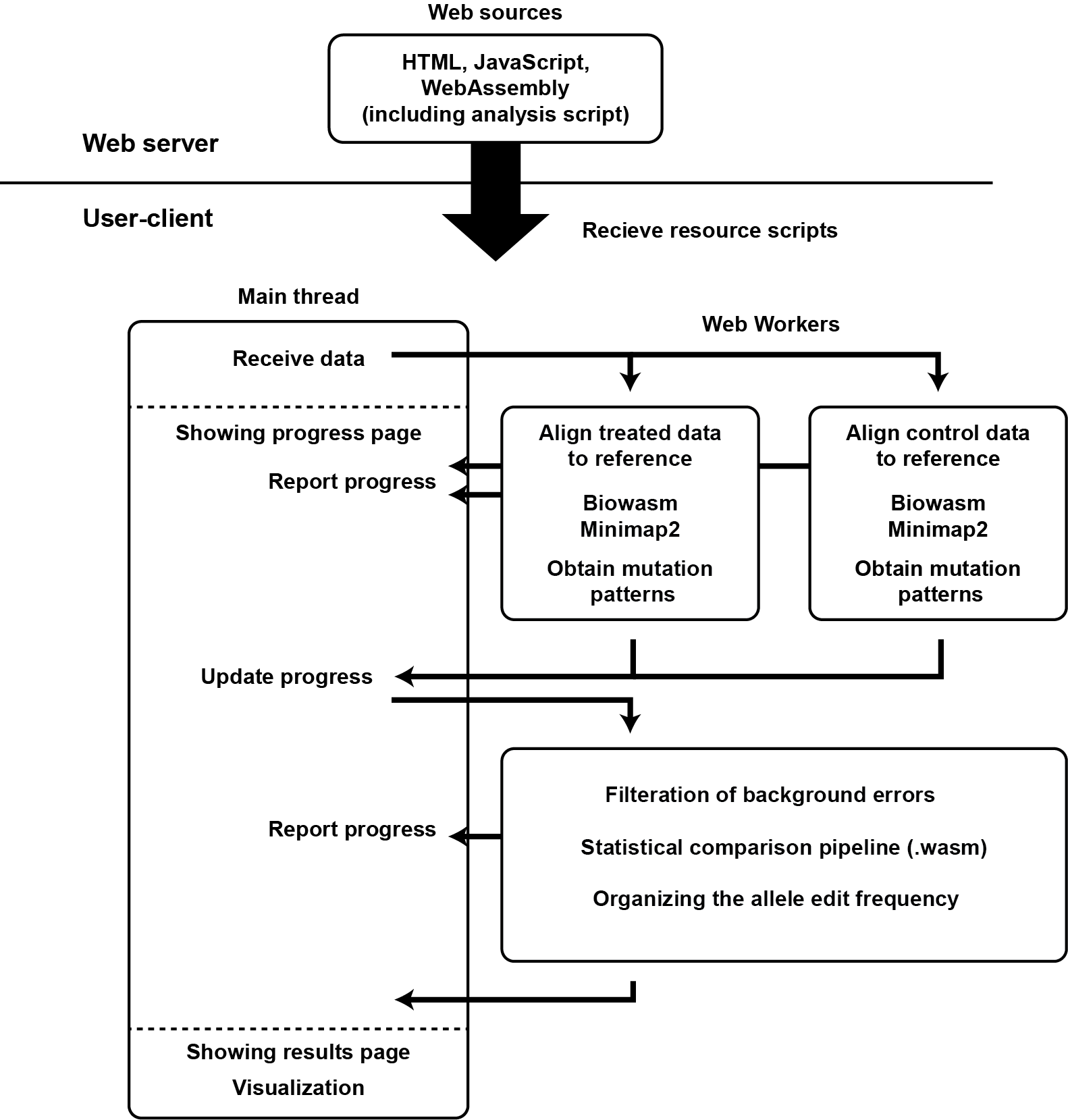

**Supplementary Figure 2. Schematic overview of the CRISPRlungo web tool.**

The HTML, JavaScript, and WebAssembly scripts required for analysis are downloaded from the web server to the user's browser. When the user inputs data, the browser's main thread verifies the input and generates two Web Workers for alignment. Each worker loads the Biowasm-based Minimap2 WebAssembly script and performs alignment while continuously reporting progress to the main thread. The main thread visualizes this progress in real time. After alignment, the workers transfer results back to the main thread and are terminated. A separate worker is then generated for background error filtering, which loads the corresponding WebAssembly script, processes the alignment results, and returns the filtered data to the main thread before termination. Once all analysis steps are complete, the main thread generates the result page and visualizes the final output within the browser.

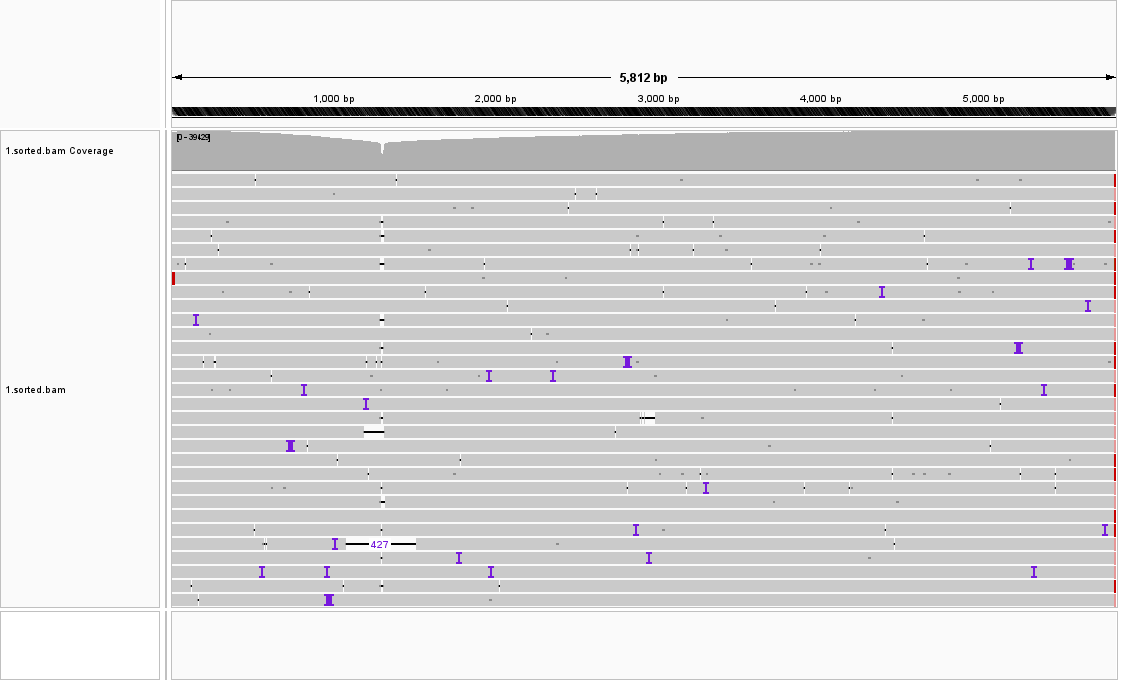

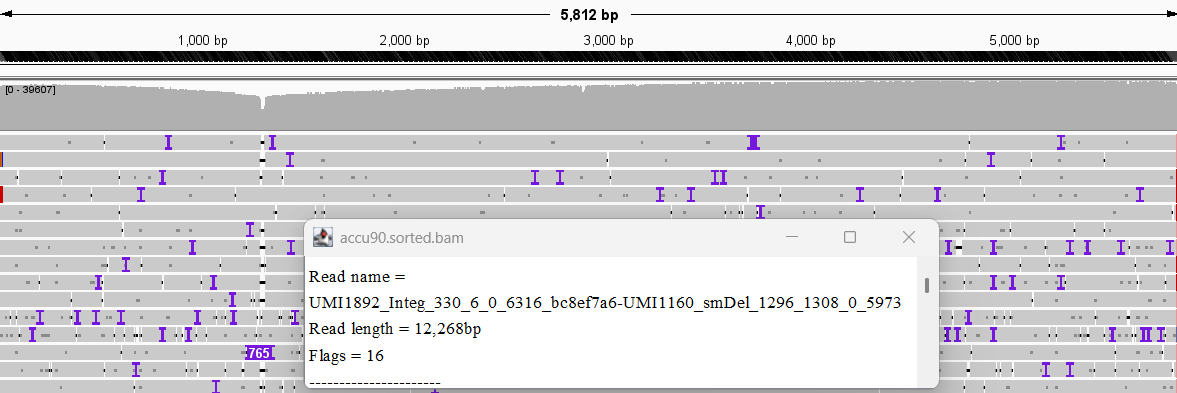

**a**

**b**

**Supplementary Figure 3. IGV alignments of long-read simulation data.**

The edited alleles were simulated using our custom pipeline, and sequencing errors were introduced with Badread at accuracies of 99% (**a**) and 90% (**b**). Information about the simulated edits was annotated in the FASTQ read headers.

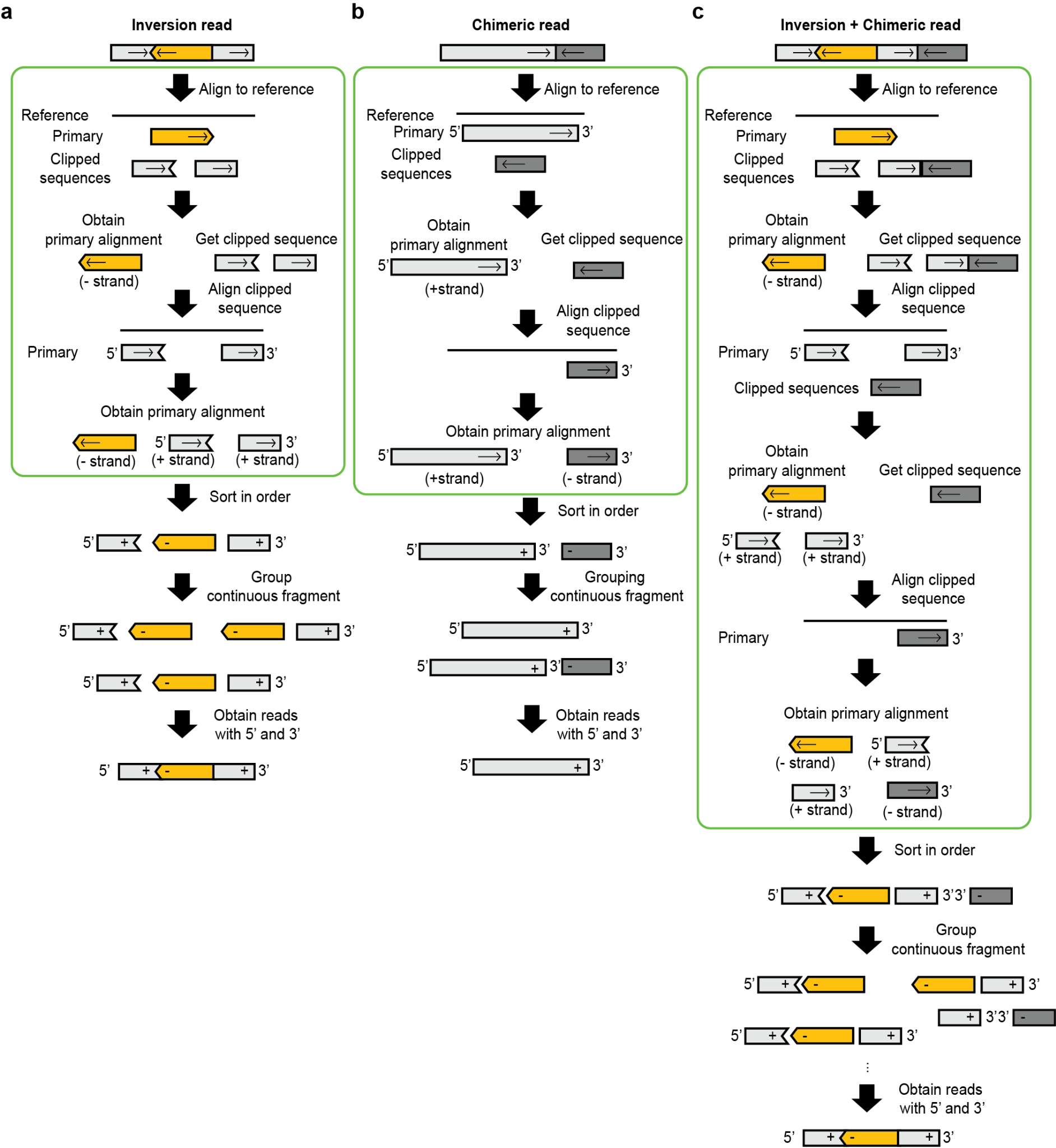

**Supplementary Figure 4. CRISPRlungo alignment pipeline.**

**a**. Example of large inversion read alignment. Orange segments represent large inversions, and arrows indicate the sequence strand relative to the reference genome. When an inversion read is aligned, the inversion region is mapped as the primary alignment. The clipped flanking sequences are extracted and re-aligned. These clipped reads are then mapped as new primary alignments. All alignment fragments are reordered according to their original sequence and combined in plausible configurations. A group of alignments with both 5' and 3' ends mapped is retained as the final alignment result. **b**. Example of a chimeric read. Dark gray segments represent different DNA molecules that were ligated during library preparation. The initial alignment yields a primary mapping of the light gray read. The clipped dark gray fragment is extracted and re-aligned. If this realignment results in a new primary alignment, it is retained. Among the alignment results, fully aligned light gray reads are preserved, while partially aligned and damaged fragments are discarded. **c**. Example of a read containing both an inversion and a chimeric fragment. The alignment process follows the same steps as in **a**. The dark gray chimeric fragment remains clipped after the inversion alignment and is re-aligned iteratively until no clipped segments remain. As with the previous cases, a plausible combination of fragments is assembled to obtain an inversion read with both ends successfully aligned.

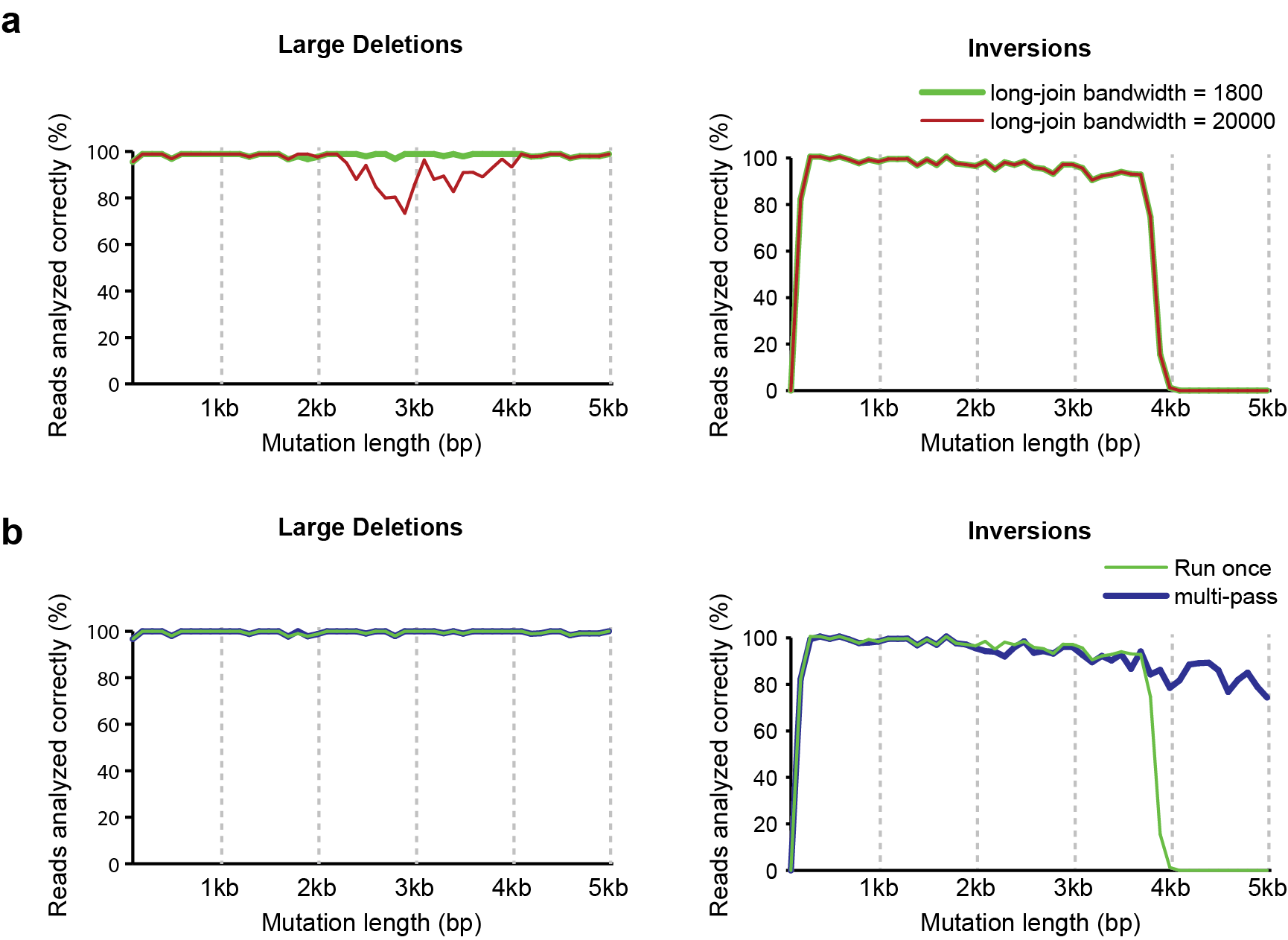

**Supplementary Figure 5. Optimization of Minimap2 alignment parameters.**

Two long-join bandwidth values were compared using simulation data for large deletions (**a**) and inversions (**b**). Run-once and multi-pass alignments were compared using simulation data for large deletions (**c**) and inversions (**d**). "Run once" (light green) indicates alignment using a single run of Minimap2. "Multi-pass" refers to the improved alignment pipeline implemented in CRISPRlungo. The line plots show the frequency of correctly analyzed reads at each mutation length.

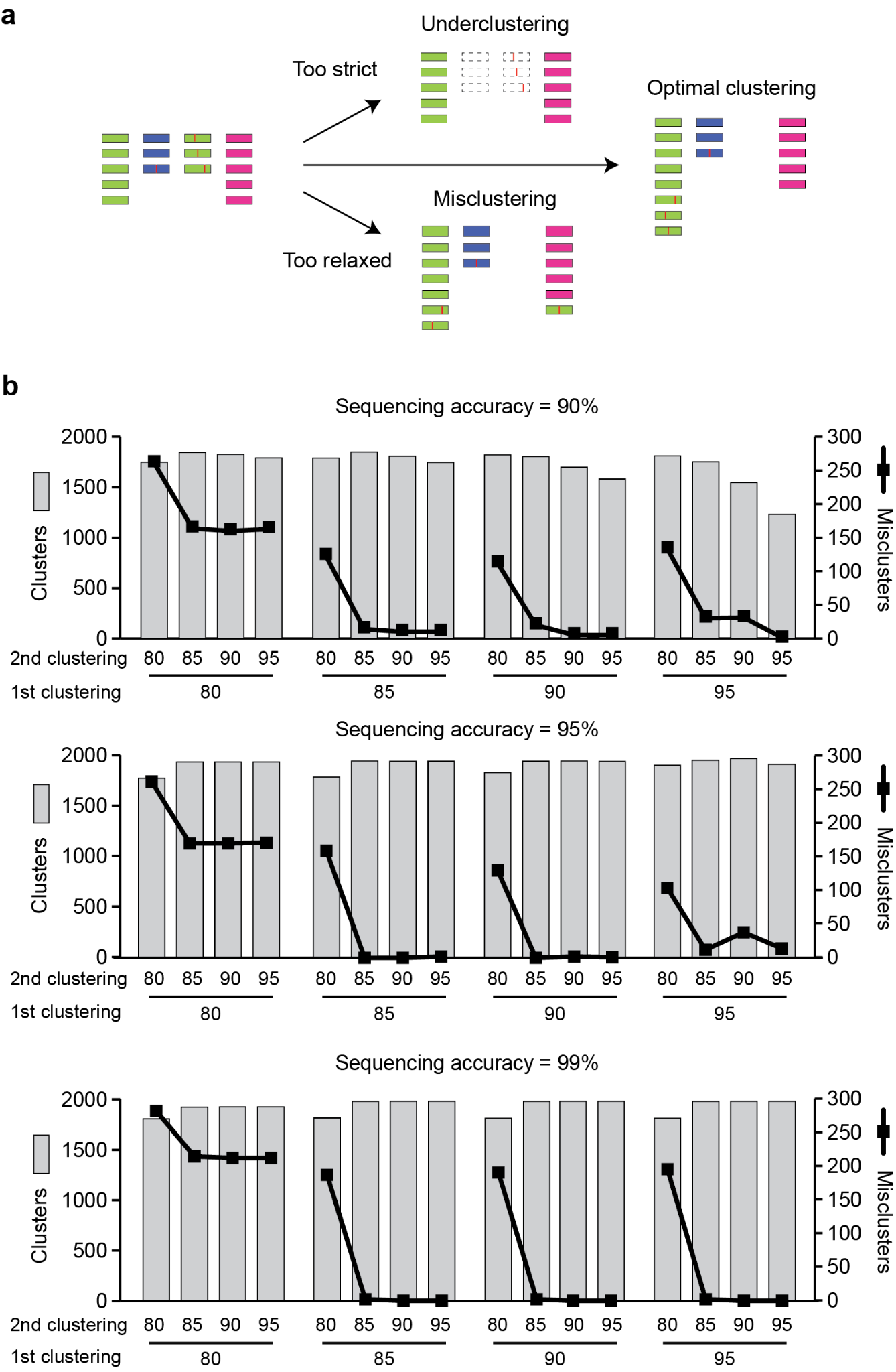

**Supplementary Figure 6. Optimization of UMI clustering.**

**a**. Schematic showing the effect of UMI clustering stringency on analysis outcomes. **b**. Results of clustering option optimization. All possible identity combinations from two VSEARCH runs were tested. The bar plots represent the total number of UMI clusters, while the line plots indicate the number of misclusters. The analysis was conducted at sequencing accuracies of 90% (top), 95% (middle), and 99% (bottom).

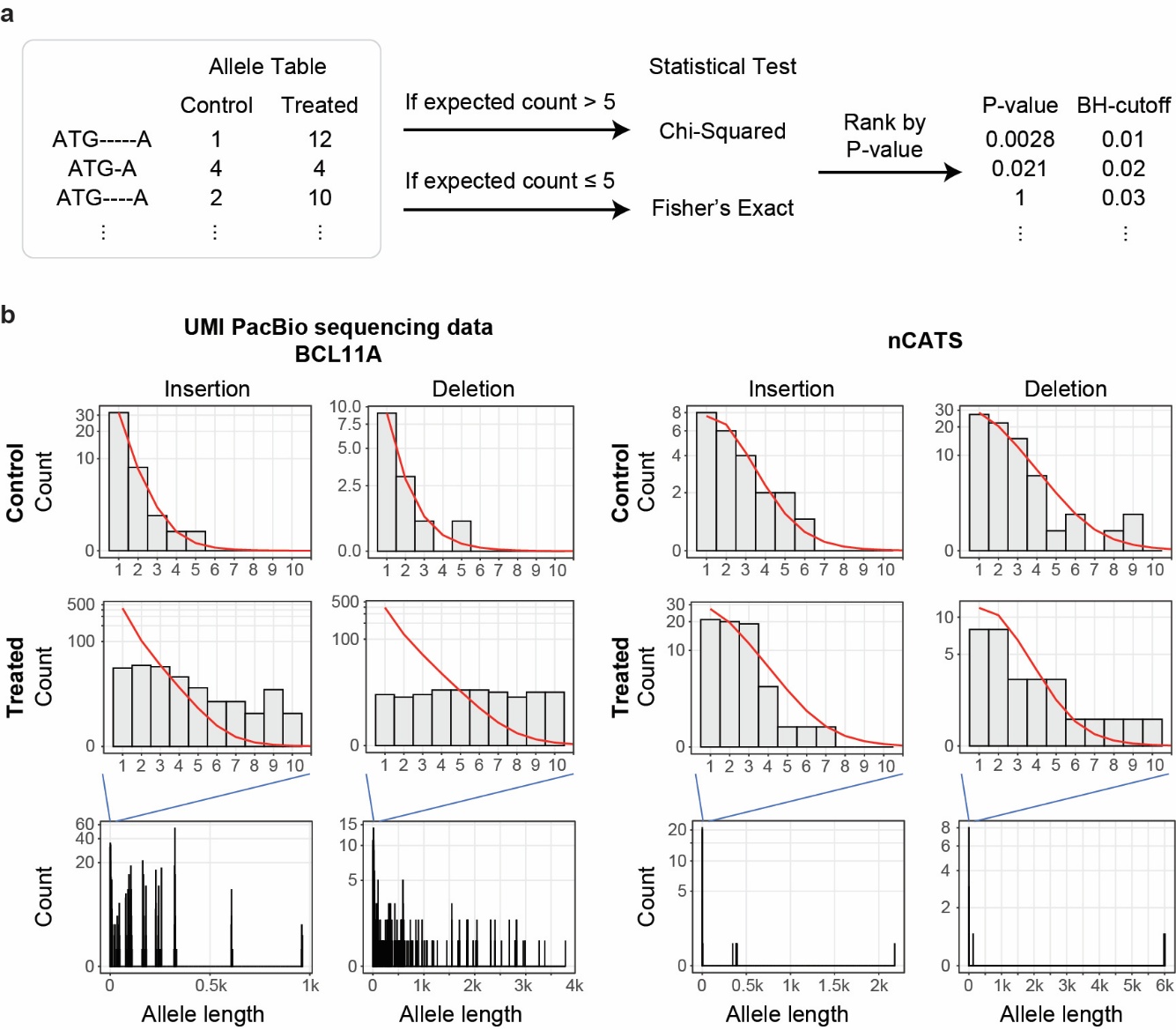

**Supplementary Figure 7. Calibration of statistical tests for error filtering.**

**a.** Schematic showing the calibration of statistical tests to filter out false positive events. **b**. Distribution of insertion and deletion allele lengths in treated and control samples. Histograms show the distributions for insertions and deletions within the default editing window from UMI PacBio sequencing of the *BCL11A* locus (left) and the nCATS dataset (right). The red curves represent the shifted negative binomial fits. Events longer than 10 bp are rare in control data (1/158), supporting a conservative cutoff of >10 bp to define significant large indels.

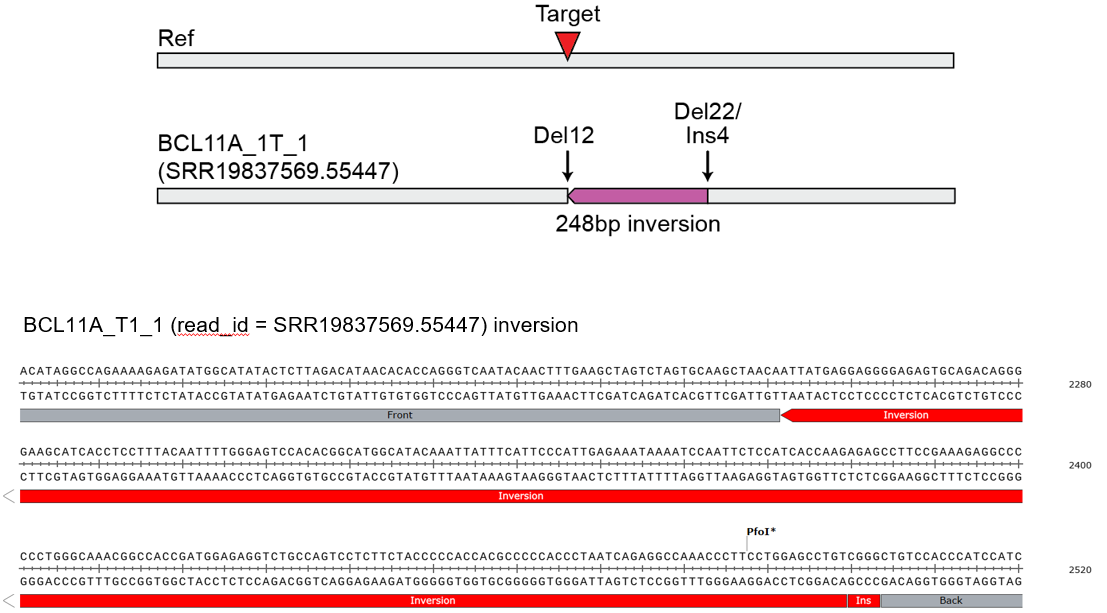

**Supplementary Figure 8. Detection of a previously unreported inversion in published dataset using CRISPRlungo.**

The top panel visualizes the inversion (purple) and accompanying indels relative to the reference sequence. The bottom panel shows the actual read sequence from the FASTQ file aligned and visualized accordingly.

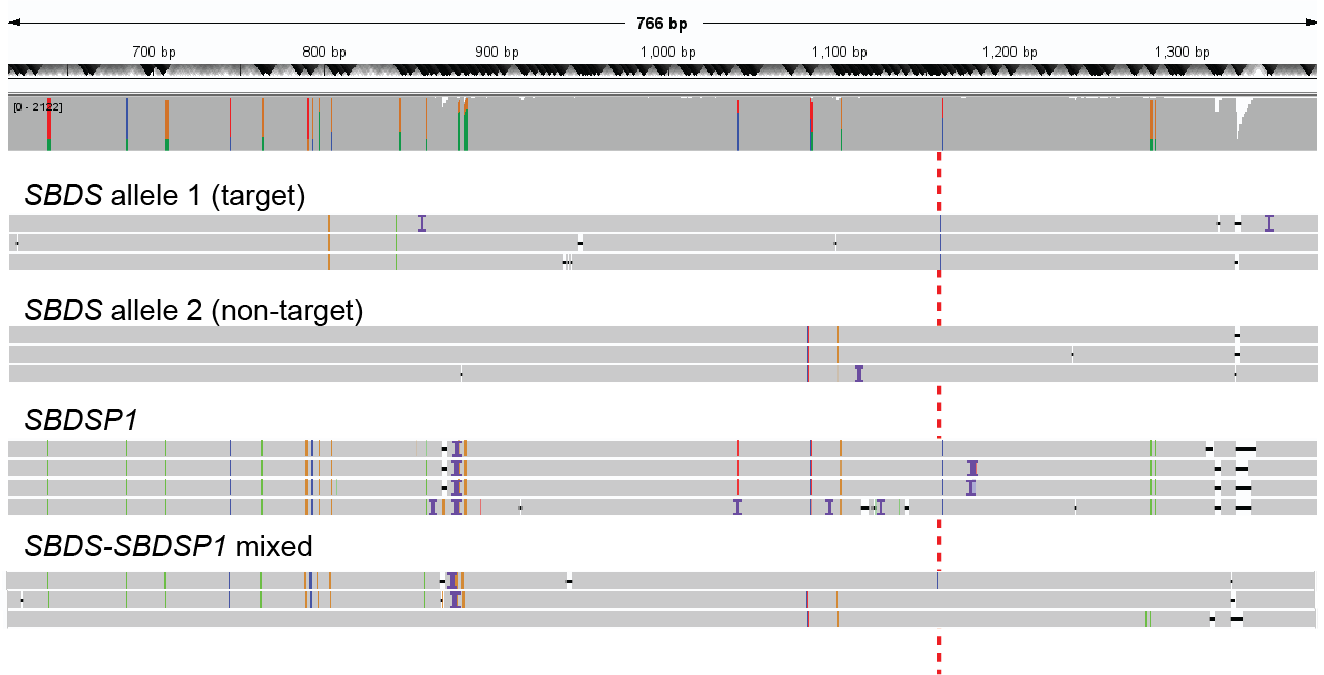

**Supplementary Figure 9. Presence of contaminating *SBDSP1* reads in IGV visualization of the mock (unedited) *SBDS* sample.**

The mock sample was aligned to the SBDS reference and visualized in IGV. In this region, SBDSP1 exhibits multiple substitutions (vertical colored lines) relative to SBDS at positions 700~900 bp and 1,283~1,285 bp of the reference. Additionally, chimeric reads containing only one side of the SBDSP1-specific mutations were detected.

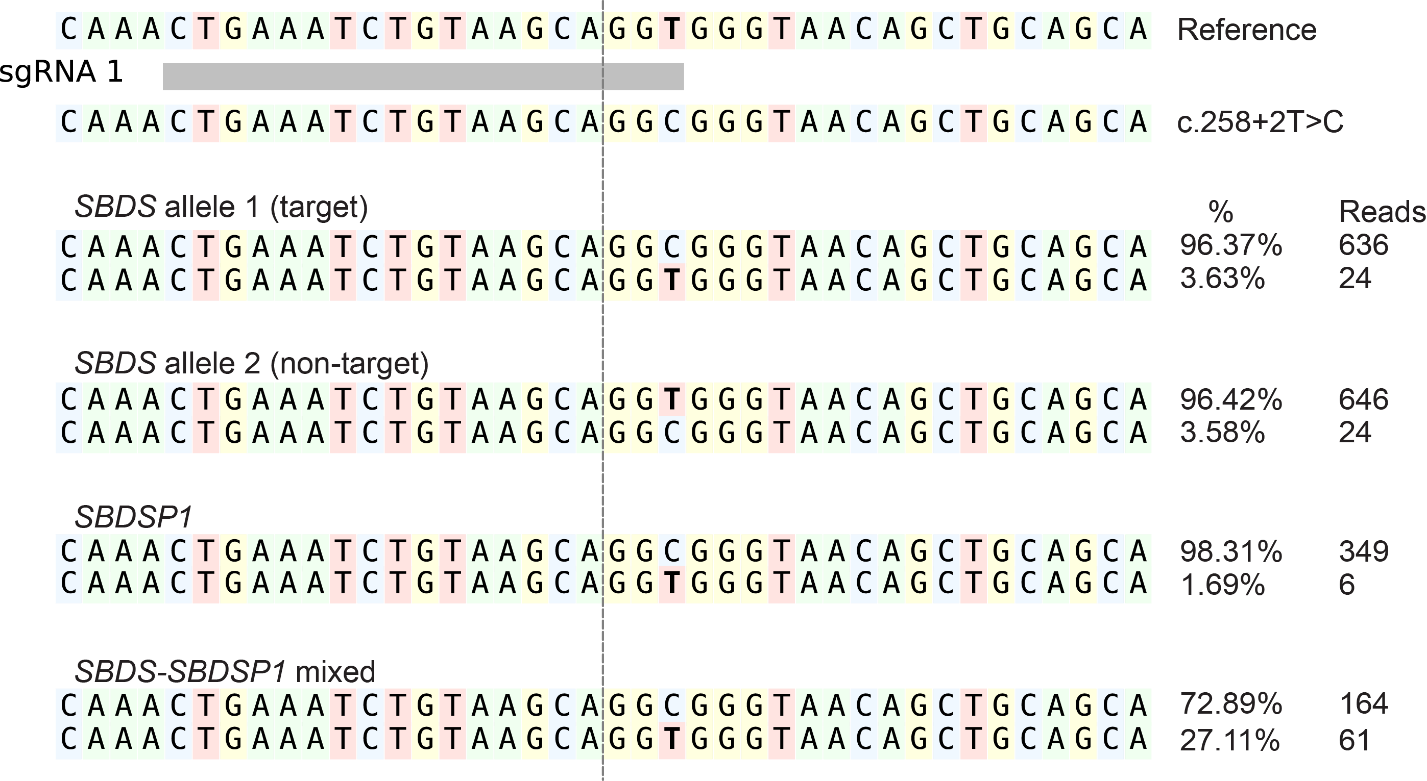

**Supplementary Figure 10. Allele-specific edit analysis of the mock (unedited) sample.**

Red boxes indicate insertions, horizontal lines represent deletions, and dotted lines indicate the cleavage sites.

**Supplementary Note 1: CRISPRlungo command line instructions and output**

**CRISPRlungo**

**CRISPRlungo** is a software pipeline designed to analyze genome editing outcomes using long-read sequencing data. It supports multiple CRISPR platforms, including base editors and prime editors, and is compatible with various sequencing methods such as amplicon sequencing, UMI-tagged long-read sequencing, and nCATS.

**Pipeline Overview**

1. Align sequencing reads to the reference genome, filter out low-quality reads, and remove chimeric reads.
2. If UMIs are used, cluster UMI-tagged reads and generate consensus sequences.
3. If control samples are provided, perform background error filtering using statistical analysis.
4. Quantify small deletions (<100bp), small insertions (<20bp), large deletions (>200bp), large insertions (>20bp), and inversions. (Because the definition of the boundary between small and large mutations differs across their conditions, options were provided to adjust this threshold.)
5. Use the submodule CRISPRlungoAllele to classify allele groups and identify PCR-induced chimeric reads.

**What can CRISPRlungo do?**

**CRISPRlungo** enables comprehensive analysis of long-read sequencing data from genome editing experiments. Its key features include:

- Filtering of low-quality reads
- Identification and removal of chimeric reads generated during library preparation
- Alignment using an optimized pipeline to detect large structural variants and inversions
- UMI extraction, clustering, and consensus read generation (if UMIs are present)
- Background error estimation and filtering using statistical methods (if control data is available)
- Quantification of small insertions/deletions (indels), large indels, inversions, and sequence integrations
- Detection and quantification of intended mutations when a reference for the edited sequence is provided
- Visualization of indel size and position distributions
- Visualization of substitution patterns and their positions
- Visualization of allele frequency and edit spectrum

**Installation**

**CRISPRlungo** can be installed by downloading from GitHub and below commands.

git clone https://github.com/pinellolab/CRISPRlungo

cd CRISPRlungo

conda env create -n {env_name} -f environment.yml # can use mamba instead of conda

pip install -e .

CRISPRlungo -h

**CRISPRlungo usage**

**CRISPRlungo** has 4 options to use.

*Using default option*:

CRISPRlungo {reference.FASTA} {sequencing_result.FASTQ} {Output directory} {target sequence}

*Using background filtering option*:

CRISPRlungo {reference.FASTA} {sequencing_result.FASTQ} {Output directory} {target sequence} –control {sequencing_result_for_mock.FASTQ}

*Using umi filtering option*:

For UMI option the UMI context must be annotated at reference FASTA file using “(“ and “)” like below.

>PD1

GGTGCTGAAGAAAGTTGTCGGTGTCTTTGTGTTAACCGTATCGTGTAGAGACTGCGTAGGTTT(VVVVTTVVVVTTVVVVTTVVVV)TTTGGGACACCGTATGTGTT…

CRISPRlungo --umi {reference.FASTA} {sequencing_result.FASTQ} {Output directory} {target sequence}

*Using background filtering option*:

CRISPRlungo –umi {reference.FASTA} {sequencing_result.FASTQ} {Output directory} {target sequence} –control {sequencing_result_for_mock.FASTQ}

Example run with background error filter

cd data

CRISPRlungo PD1.fasta --control Nanopore_umi_Run_test_control_wo_chi.fastq Nanopore_umi_Run_test_wo_chi.fastq regular_output ggcgccctggccagtcgtct

This command generate “regular_output” folder and you can see the results at regular_output/combined_graphs.html.

Example run with background error filter

cd data

CRISPRlungo --umi PD1_umi.fasta --control Nanopore_umi_Run_test_control_wo_chi.fastq Nanopore_umi_Run_test_wo_chi.fastq umi_output ggcgccctggccagtcgtct

This command generate “umi_output” folder and you can see the results at umi _output/combined_graphs.html.

**Parameter list**

--umi: Enable this option if UMI sequencing was used.

--control: Provide a control file to perform background filtering using the control data.

--cleavage_pos: Cleavage position relative to the target sequence. [default: 16]

--additional_target: Additional target sequence to include in the analysis.

--window: The analysis window size around the cleavage site.

--whole_window_between_targets: Include the entire region between two targets in the mutation analysis window.

--induced_sequence_path: Provide a file containing the desired induced sequence to enable additional classification of specific mutations.

--integration_file: FASTA file containing sequences that may be integrated into the genome.

--merge_substitution: Treat consecutive substitutions as a single mutation event.

--min_read_cnt: Filter out reads fewer than the received value.

--min_read_freq: Filter out reads with frequencies below the received value.

--mix_tag: If multiple mutations are detected in a read, set to "False" to prioritize indels (labeling them as ins or del), or "True" to label them as Complex mutations. [default: False]

--min_mut_freq_no_control_refmut: Minimum mutation frequency threshold for classifying mutations in the absence of a control file and desired sequence.

--induced_paritial_similiarity: If a mutation pattern is not identical to the desired sequence but shows similarity above this threshold, it will be considered partially induced.

--range_both_end_region: Reads that are not aligned within this range from both ends are considered short fragments.

--align_sa_len_threshold: Minimum alignment length threshold for soft-clipped or supplementary alignments.

--p_value_threshold: Statistical p-value threshold for significance filtering.

--mut_freq_threshold: Mutation frequency threshold for filtering; increase this value for more stringent filtering.

-c or --clust_cutoff: Minimum UMI cluster size threshold.

--just_visualization: Skip consensus generation and only analyze mutations if consensus reads have already been generated using CRISPRlungo. [default: False]

--allele_plot_window: Window size used for allele plots. [default: window + 10]

--allele_plot_lines: Number of representative sequences shown per allele plot. [default: window + 10]

--show_all_between_allele: Draw all sequences spanning the region between two targets in the allele plot.

-t or --threads: Number of threads to use for parallel processing.

**Result files descriptions.**

*Combined_graphs.html* shows the organized results and graphs in a web browser.

*Input_summary.txt* displays the analysis options used in the run.

*Allele_table.txt* is a tab-separated file that shows information associated with each allele. Within the defined window, the first and second columns display the aligned reference and sequencing results, respectively. The third column shows the count of each allele, and the fourth column presents their frequency. The fifth column lists the mutations in CIGAR string format, while the sixth column indicates the position and type of each mutation.

*Edit_pattern_count.txt* reports the count and frequency of each mutation pattern. The first column lists the mutation pattern; the second and third columns show the count and frequency, respectively. The fourth column indicates integration information, the fifth column shows whether the mutation was induced, and the sixth column provides the position and length of the mutation.

*Mutation_pattern_p_values.txt* provides the p-values for each mutation pattern. The first column lists the mutation pattern, and the third column shows the corresponding p-value. Columns four through seven display the total read count and mutation count for the control and treated datasets, respectively. The eighth column indicates whether the mutation pattern was included in the final analysis.

*Mutation_summary_count.txt* summarizes the number of reads filtered during preprocessing and the read counts for each mutation pattern.

*Preprocess_count.txt* summarizes the number of reads filtered during the preprocessing process.

*Read_classification.txt* contains the mutation analysis results for each read. The first column shows the read ID, the second column indicates the mutation classification, the third column lists the mutation pattern within the defined window, the fourth column provides integration status, the fifth column specifies whether the mutation is induced, and the sixth column presents all mutations detected in the read.

An example output is Fig. S11.

**Subanalysis tool – CRISPRlungoAllele**

**CRISPRlungoAllele** performs post-analysis of CRISPRlungo results by classifying alleles into multiple groups based on mutation type.

This enables:

- Identification of mutation patterns within alleles
- Detection of similar/homologous sequences
- Estimation/removal of PCR chimeric products

**CRISPRlungoAllele** is executed using the following command:

CRISPRlungoAllele {analysis_file_dir} {custom_category_file}

analysis_file_dir: the output directory used in the original CRISPRlungo run.

custom_category_file: tab-separated text file formatted as follows:

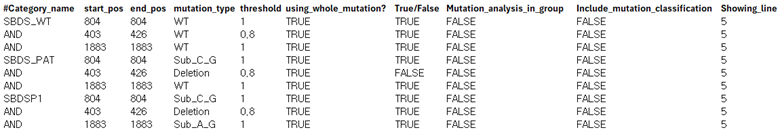

Each row in the custom_category_file defines a mutation condition for a specific category.

·	The first column specifies the name of the category. If multiple mutation conditions belong to the same category, enter "AND" in the first column of subsequent rows to group them.

·	The second and third columns indicate the start and end positions of the mutation.

·	The fourth column defines the mutation type: "WT", "Sub", or "Deletion". For "Sub", the reference and mutated nucleotides should be written as ref_nt_mut_nt (e.g., A_G).

·	The fifth column sets the threshold for how much of the specified region must meet the condition (e.g., 403/426/WT/0.8 means that the region from 403 to 426 bp must remain at least 80% unmutated to satisfy the condition).

·	The sixth column indicates whether to apply the condition only to mutations within the defined analysis window. Use "TRUE" to include mutations outside the window.

·	The seventh column determines whether the condition must be met (TRUE) or not met (FALSE) to classify the read into the category.

·	The eighth column specifies whether to calculate mutation frequency within the group (TRUE) or across all reads.

·	The tenth column sets how many top alleles from this category will be shown in the final allele plot.

**Example:** CRISPRlungoAllele regular_output customized_mutation.txt

**CRISPRlungoAllele** accepts the following four options:

--min_read_cnt: Filter out reads fewer than the received value.

--min_read_freq: Filter out reads with frequencies below the received value.

--allele_plot_window Window size used for allele plots. [default: window + 10]

--show_all_between_allele Draw all sequences spanning the region between two targets in the allele plot.

**CRISPRlungoAllele result files descriptions.**

**CRISPRlungoAllele** creates a “custom_results” folder within the output directory, which contains the analysis results for each defined category. Inside the “custom_results” folder, the following files are generated: “allele_plot.png”, “custom_mutation_allele_plot_input.txt”, and “custom_mutation_pattern_count.txt”. Each category-specific subfolder includes the following files: “deletion_and_insertions_per_position.png”, “mutation_pattern_count.txt”, “Mutation_pie_chart.png”, “mutation_summary_count.txt”, “pattern_pie_chart.png”, “percent_of_alleles_pie_chart.png”, and “read_classification.txt”. These files follow the same format as those generated by the original CRISPRlungo output.

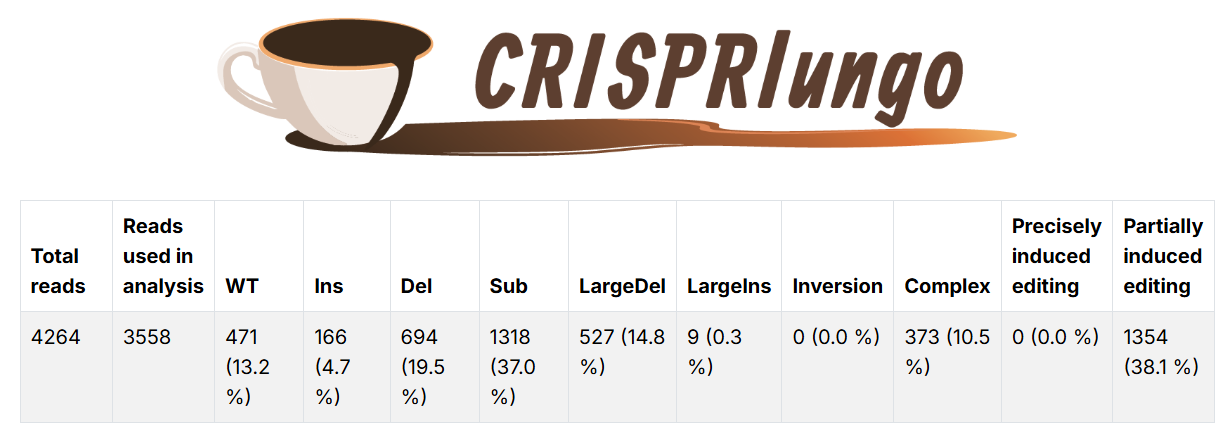

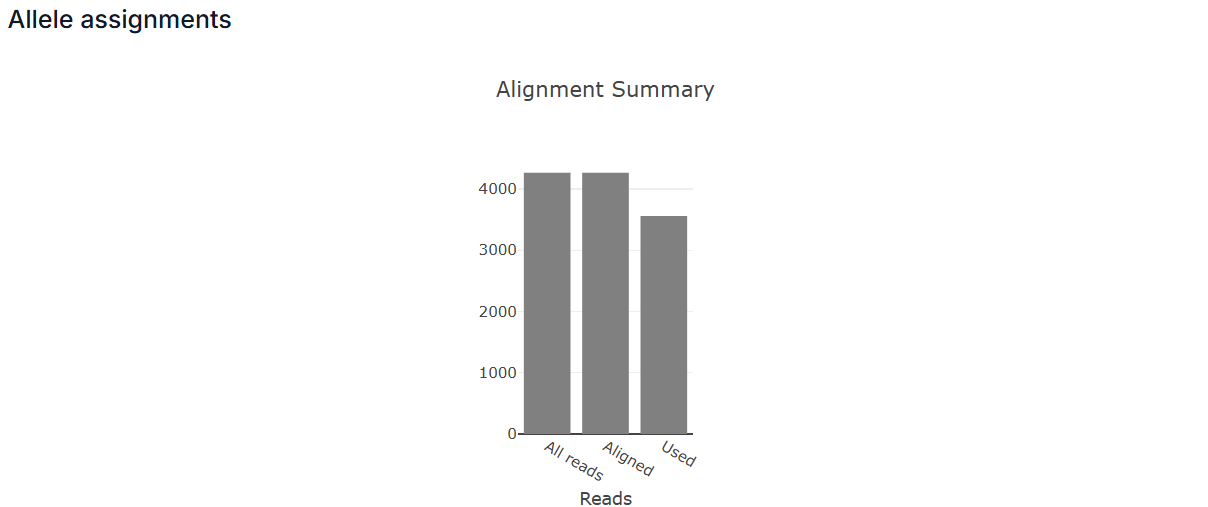

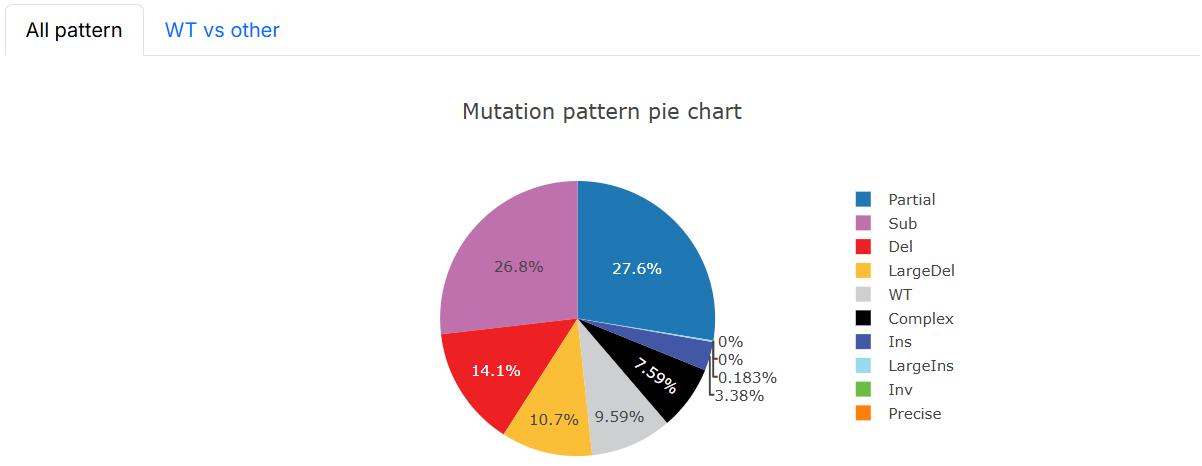

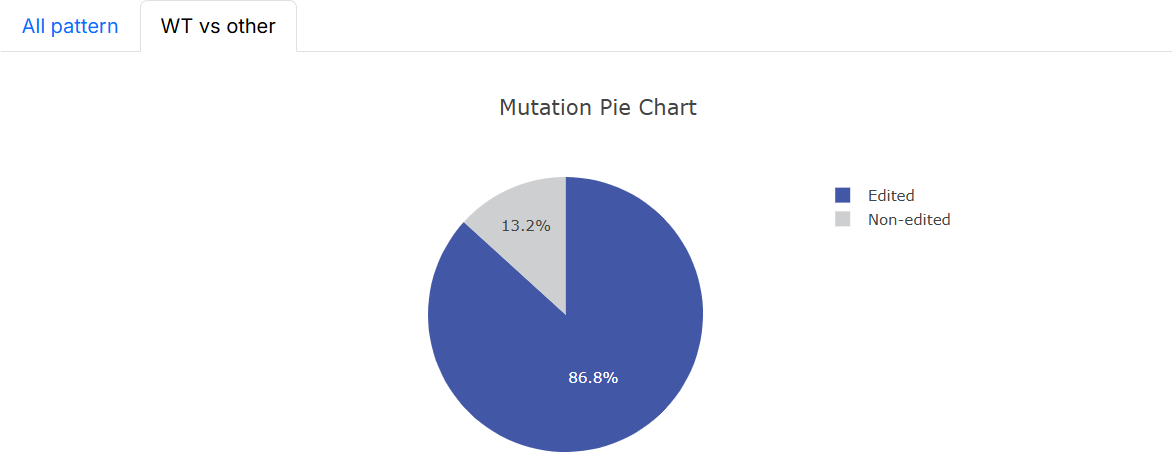

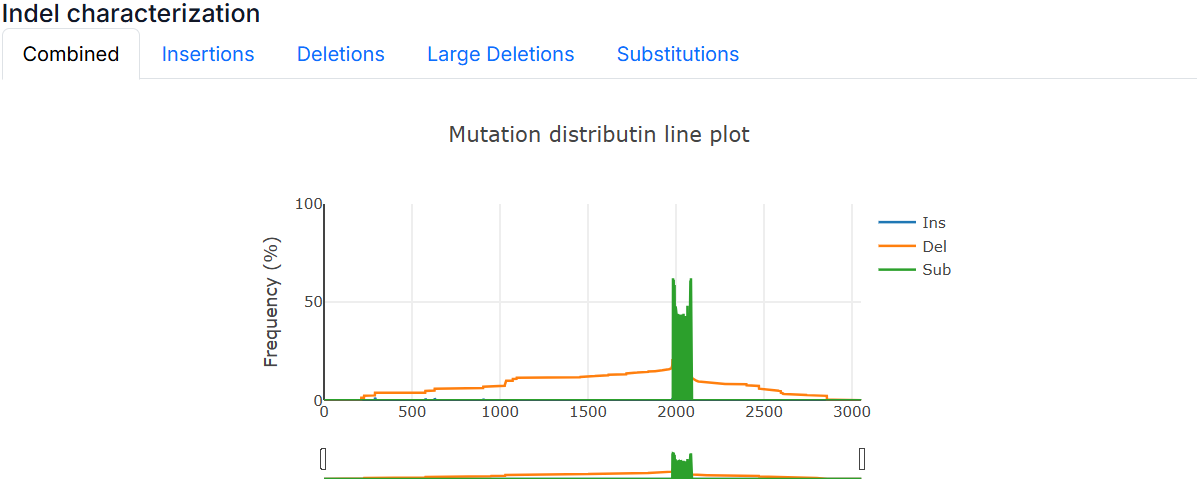

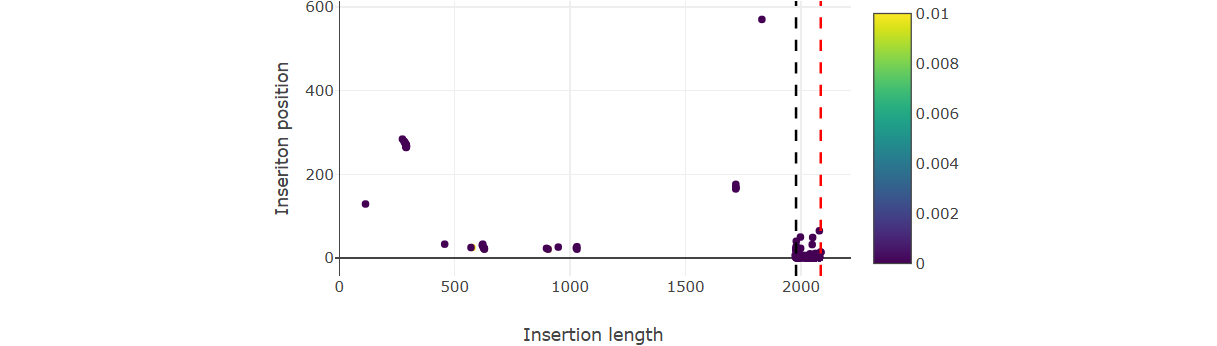

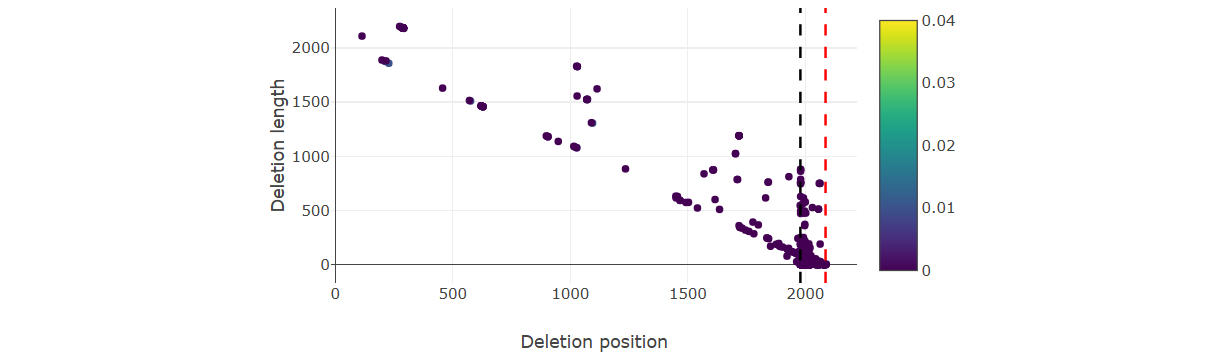

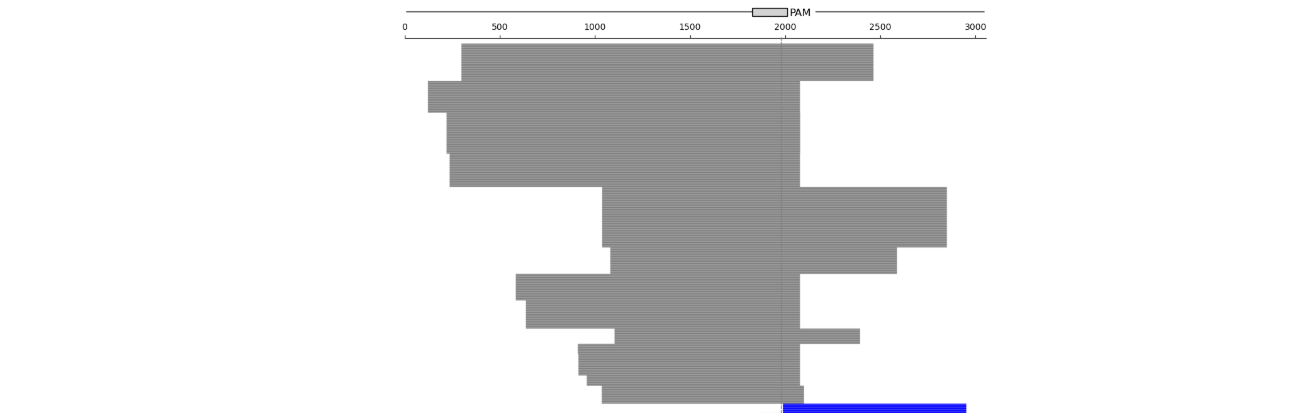

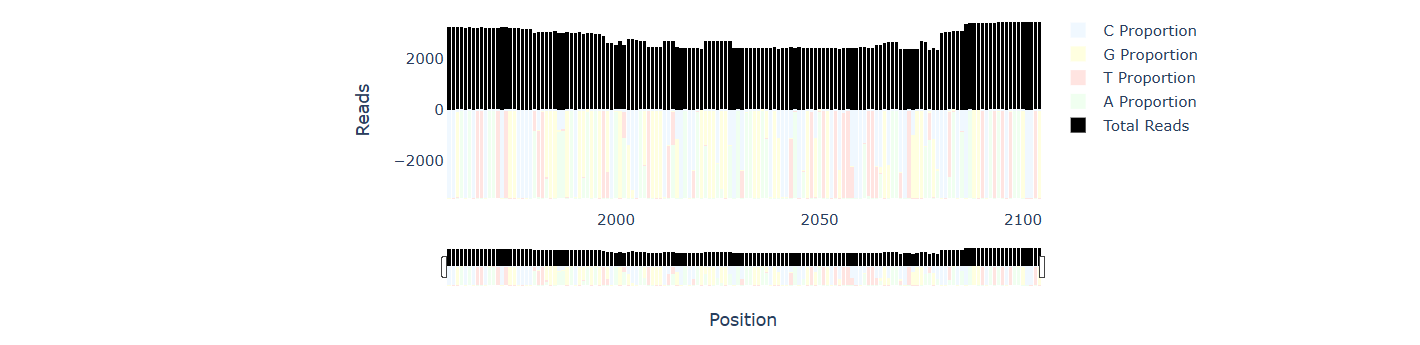

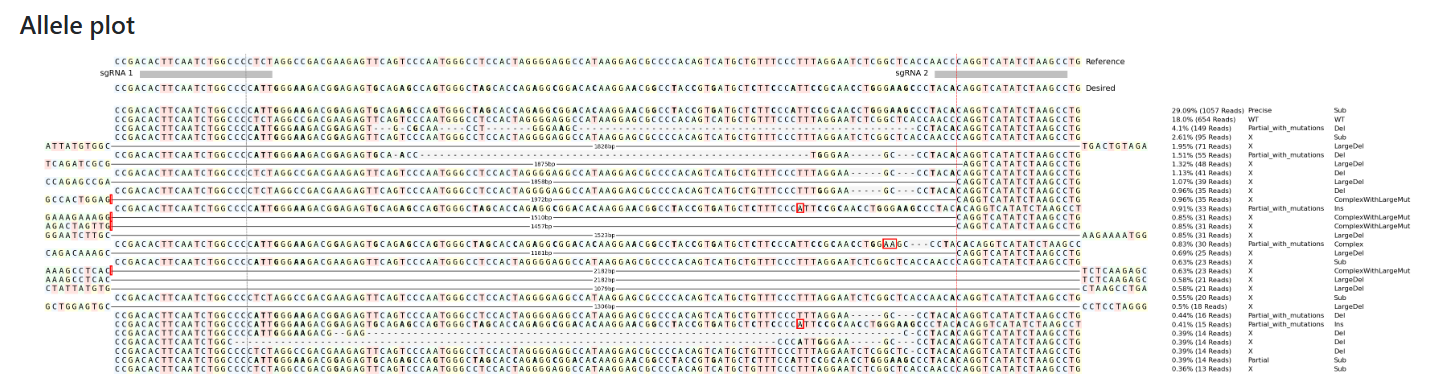

**Supplementary Figure 11. CRISPRlungo output**

CRISPRlungo output includes reports of alignment, edit frequency, mutation position, length, and allele patterns.
